# Syncing Minds for Learning: Student-Teacher Inter-brain Coupling Forecast Cross-semester Academic Fluctuation in Real-World Classrooms

**DOI:** 10.1101/2024.05.08.593270

**Authors:** Xiaomeng Xu, Dan Zhang, Yu Zhang

**Affiliations:** School of Education, Tsinghua University, Beijing, China, 100084; Department of Psychological and Cognitive Sciences, Tsinghua University, Beijing, China, 100084; Tsinghua Laboratory of Brain and Intelligence, Tsinghua University, Beijing, China, 100084

**Keywords:** student-teacher inter-brain coupling, hyperscanning, causal relationship, longitudinal study, authentic classroom

## Abstract

Student performance exhibits dynamic fluctuates throughout the learning process, which reflects deeper educational myths and impacts educational quality and equity. Although teachers are believed to play a pivotal role in shaping these changes, the mechanisms through which student-teacher interactions drive performance variations across individual students remain unclear. The present study investigates student-teacher inter-brain coupling in authentic classroom environments through longitudinal neurophysiological recordings spanning three academic semesters. Using wearable hyperscanning technology, a total of 4,175 longitudinal EEG recordings were collected from 107 junior and senior students across 393 regular Chinese and math classes. While static models revealed a negative association between student-teacher inter-brain coupling and math performance—likely reflecting teacher selection effects—dynamic analyses showed that increased coupling, particularly in the high-beta frequency band, causally predicted academic improvement in both subjects. These findings highlight the functional significance of inter-brain coupling in real-world classroom learning, providing robust ecological validity and supporting its potential as a measurable marker of effective pedagogy.

**Significance Statement:** This study is the first to provide causal evidence, based on real-world classroom data, that enhanced student-teacher inter-brain coupling improved academic performance. Leveraging 4,175 longitudinal EEG recordings from 107 students across three semesters, we show that higher coupling levels predict better learning outcomes over time. Capturing neural dynamics in authentic classrooms, this research offers high ecological validity and strong potential for practical application and large-scale educational implementation.

## Introduction

Across diverse educational settings and cultures, student performance is inherently dynamic, marked by differentiation and fluctuation over time. These fluctuations reflect deeper educational myths and raise critical questions about the nature of educational quality and equity. As learners progress, their learning trajectories diverge, shaped by a complex interplay of individual, social, and instructional factors. Central to this process is classroom instruction—particularly the quality of student-teacher interaction—which serves as the cornerstone of formal learning. These interactions not only facilitate transmit knowledge but also shape student engagement, motivation, and cognitive development (Hamre & Pianta, 2007; Holper et al., 2013; Pianta et al., 2012; Saha & Dworkin, 2009). Yet, in everyday practice, educators often face difficulties in discerning which students are keeping pace, which are falling behind, and how their own instructional practices contribute to students’ learning outcomes. Addressing this gap requires deeper insights into how student-teacher interactions drive dynamic trajectories, especially among students with varying learning profiles. Such insights are essential for developing adaptive, evidence-based pedagogical strategies that foster equitable learning environments and empower all students to thrive.

However, effectively measuring the quality of student-teacher interactions presents significant methodological challenges. These interactions are inherently complex, encompassing both observable behaviors, such as questioning and responding, and implicit cognitive and affective exchanges (Ellis, 1993; Ellis, 2005; Hamre & Pianta, 2005). For instance, experienced teachers can intuitively tailor their instruction through such interactions to align with students’ zones of proximal development, crafting learning experiences that resonate with students’ current abilities and potential for growth (Shabani et al., 2010; Vygotsky, 1978). Therefore, dynamically assessing the quality of student-teacher interactions is crucial for gaining a deeper understanding of students’ learning outcomes. Yet, objectively and accurately quantifying these interactions remains elusive. Conventional assessments, such as self-reported questionnaires, are limited by subjective biases, social desirability effects and potential measurement errors (Assor & Connell, 1992; Miller, 2011; Nederhof, 1985; Pace, 1985).

Recent advances in EEG-based hyperscanning offer a transformative approach to study student-teacher interactions in ecologically valid classroom settings. This technique captures inter-brain coupling—a neural biomarker of knowledge transmission and inter-personal interaction.—with high temporal resolution and minimal invasiveness (Feng et al., 2025; Meshulam et al., 2021; Zheng et al., 2018; Zhu et al., 2022). While prior lab-based studies have linked inter-brain coupling to learning outcomes (Meshulam et al., 2021), its role in authentic classrooms remains underexplored, particularly across longitudinal timescales. Wearable EEG systems now enable sustained, real-world data collection, overcoming limitations of bulky neuroimaging tools (e.g., fMRI/fNIRS) and addressing the dynamic, bidirectional nature of classroom exchanges. Challenges such as confounding factors (e.g., prior knowledge, teacher bias) can be addressed through longitudinal econometric modeling, which isolates the predictive power of inter-brain coupling dynamics on academic progress (Angrist & Pischke, 2009).

To unpack this relationship, we adopted both static and dynamic modeling approaches, offering distinct yet complementary analytical perspectives. The **static model** assesses how momentary coupling relates to concurrent academic outcomes, capturing more superficial, state-like associations. However, such associations are prone to endogeneity, especially due to complex, unobserved confounders in real classrooms—such as differences in students’ prior knowledge, cognitive abilities, motivation, learning strategies, and teachers’ self-selection in allocating attention based on student needs or performance (Angrist & Pischke, 2009). For instance, teachers may attune more closely to lower-performing students, leading to higher coupling values negatively associated with outcomes. To address these limitations, we developed a **dynamic modeling framework** leveraging longitudinal data to estimate how fluctuations in student-teacher coupling across semesters forecast changes in academic performance, highlighting the temporal and potentially causal role of neural coupling. This difference-based approach mitigates biases from time-invariant unobserved variables and teacher self-selection (Gandolfo, 1971; Liker et al., 1985; Tuma, 1984). By including baseline coupling as a covariate—given its association with later changes in both coupling and performance—we further reduce omitted-variable bias and improve internal validity. Together, the static and dynamic models offer a more robust, multidimensional understanding of how neural coupling between students and teachers relates to learning in naturalistic classrooms..

The study is based on a longitudinal investigation involving 107 students in junior and senior high school. A total of 393 classroom sessions were recorded across Chinese and math subjects in authentic learning environments. Neurophysiological data were collected using portable wearable EEG devices from both teachers and students over the course of three academic semesters, encompassing six waves of valid longitudinal data. Given the distinct functional significance of different EEG frequency bands, separate analyses were conducted for each band. For each participant, we calculated the average levels of student-teacher inter-brain coupling, as well as standardized scores representing students’ learning outcomes, for each half-semester (i.e., one *period*). To infer the predictive role of inter-brain coupling, we employed a dynamic model based on longitudinal changes and a static model based on concurrent associations. The study aims to uncover the causal relationship through which student-teacher inter-brain coupling shapes academic development over time, offering new neurophysiological insights into effective teaching-learning alignment in ecologically valid classroom contexts.

## Results

In this study, 107 students (60 males, aged 12–15) in their first year of junior and senior secondary school and eight teachers (four teaching Chinese and four teaching math) participated in the study. EEG recordings were collected during Chinese and math classes over three semesters (1.5 years in total), with one recording week per month capturing 1–2 daily sessions (40 minutes each). In total, 393 class sessions were recorded, including 182 Chinese classes (88 junior, 94 senior) and 211 math classes (113 junior, 98 senior). Prior to each session, students were equipped with EEG headbands, participated in usual class activities (Fig. 1a, b, c).

**Fig. 1.**
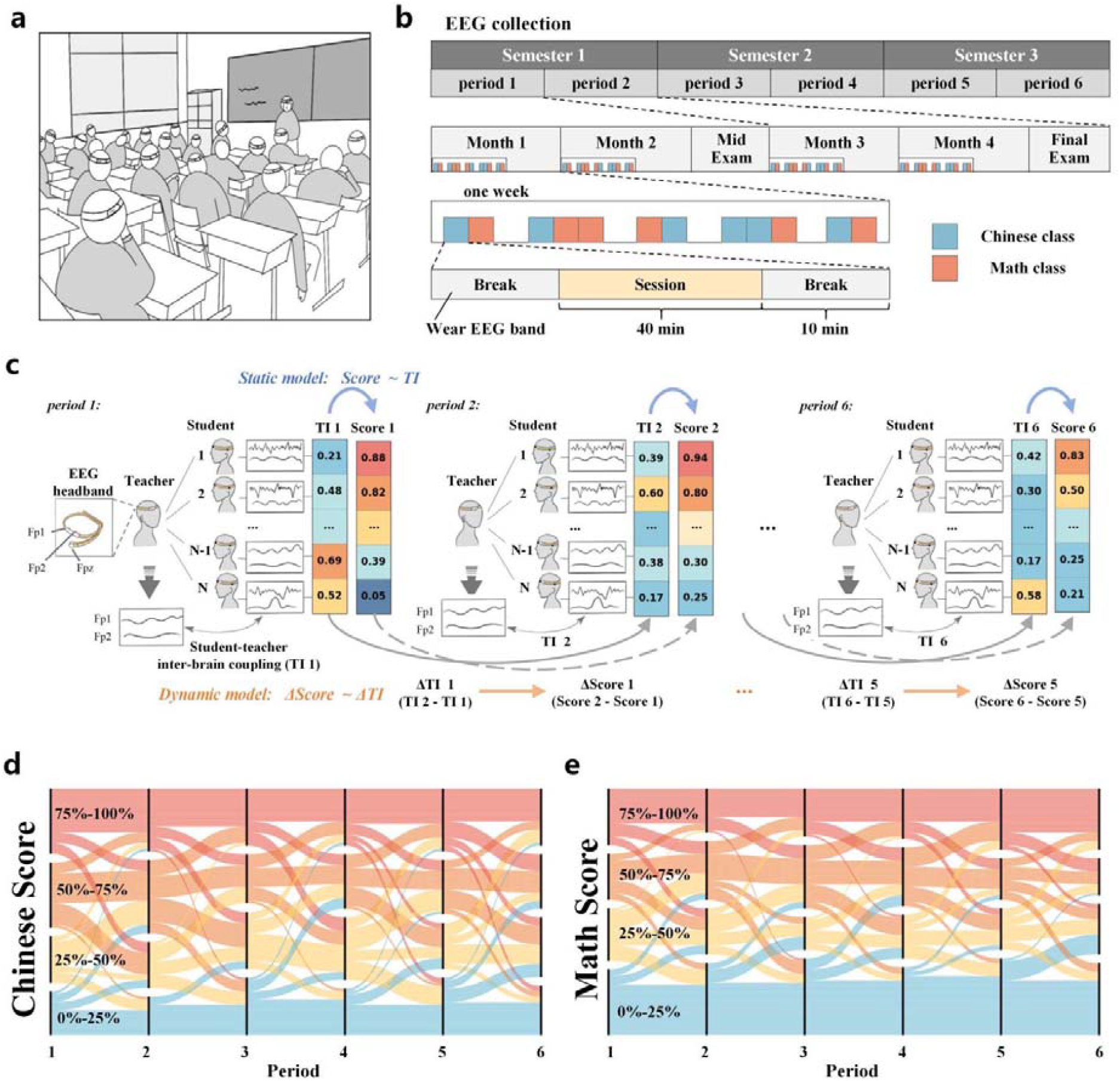
Overview of the data collection and modeling framework. **a**, Schematic of EEG data acquisition in authentic classroom. **b**, Workflow of the data collection process across three semesters. **c**, Schematic illustration of static and dynamic model construction. From left to right are the teacher–student TI values across successive periods. **d**, Variation in Chinese scores across different periods. The x-axis represents successive time points from left to right. **e**, Variation in math scores across different periods.

Students’ learning outcomes were assessed through half-semester examinations, with scores provided by the school administration. Each half-semester was operationally defined as a single period. Scores were then normalized against the entire student cohort for further analysis, with higher normalized scores indicating better performance. Fig. 1d and 1e illustrate the temporal changes in Chinese and math scores, respectively. Students were stratified into quartiles based on their grade rankings: 0-25%, 25-50%, 50-75%, and 75-100%, with the 75-100% range representing the top 25%. Notable fluctuations in student rankings were observed across performance categories over time.

The research employed the total interdependence (TI) value to quantify the student-teacher inter-brain coupling, conducting analyses across multiple frequency bands (Chen et al., 2023; Dikker et al., 2017; Xu et al., 2024). For each student, the student-teacher TI value was calculated between that student and their respective teacher during each class session and then averaged and standardized for each period. The differences in TI values and scores were calculated between two consecutive periods, and this analysis was extended across various frequency bands including broadband (1-40 Hz), delta (1-4 Hz), theta (4-8 Hz), alpha (8-13 Hz), low-beta (13-18 Hz) and high-beta (18-30 Hz) frequency bands. To explore the potential causal relationship between student-teacher inter-brain coupling and learning outcomes, the dynamic and static models were constructed to capture the relationship between student-teacher TI and test scores across periods (Fig. 1c). The dynamic model captured the relationship between changes in TI (ΔTI) and changes in test scores (ΔScore) across consecutive periods, while the static model examined the association between TI and test scores within each period.

### Changes in student-teacher inter-brain coupling forecast students’ academic improvement

The study first employed a dynamic model to investigate whether increases in student-teacher inter-brain coupling could forecast improvements in student performance. The model used the changes in learning outcomes (Δscores) between the two adjacent *periods* as the outcome variable, with the baseline student-teacher coupling values (TI values) and changes in coupling values (ΔTI values) as predictors. A Bayesian linear mixed-effects (LME) model was employed, which allowed us to account for the nested structure of the data by controlling for repeated measures across students and semesters. The Bayesian approach is well-suited for small samples and enables distribution-free inference, offering a flexible framework for modeling complex neuro data (Haug, 2012; Kruschke, 2013).

The estimation of the dynamic model was conducted separately for the full student sample, the junior student subsample, and the senior student subsample. Results for the full sample are presented in Fig. 2, while the corresponding results for the junior and senior students subsamples are provided in the *Supplementary Information* (see *Fig. S1*). After controlling the baseline TI values, the analysis showed that ΔTI values significantly predict the improvement in both Chinese (Fig. 2a) and math scores (Fig. 2b). For both Chinese and math, increases in ΔTI within the high-beta frequency band were positively associated with improvements in student scores across the full sample (Chinese: posterior mean = 0.256, 95% HPDI [0.003, 0.516]; math: posterior mean = 0.252, 95% HPDI [0.036, 0.48]). The results of the fitted value are depicted in Fig. 2c (Chinese) and Fig. 2d (math). The horizontal axis represents the predicted Δscores based on TI and ΔTI values across different frequency bands, while the vertical axis shows the actual Δscores. The regression coefficient close to 1 indicates a strong model fit and high predictive accuracy. These findings suggest that more pronounced increases in student-teacher TI values can forecast greater academic improvements.

**Fig. 2.**
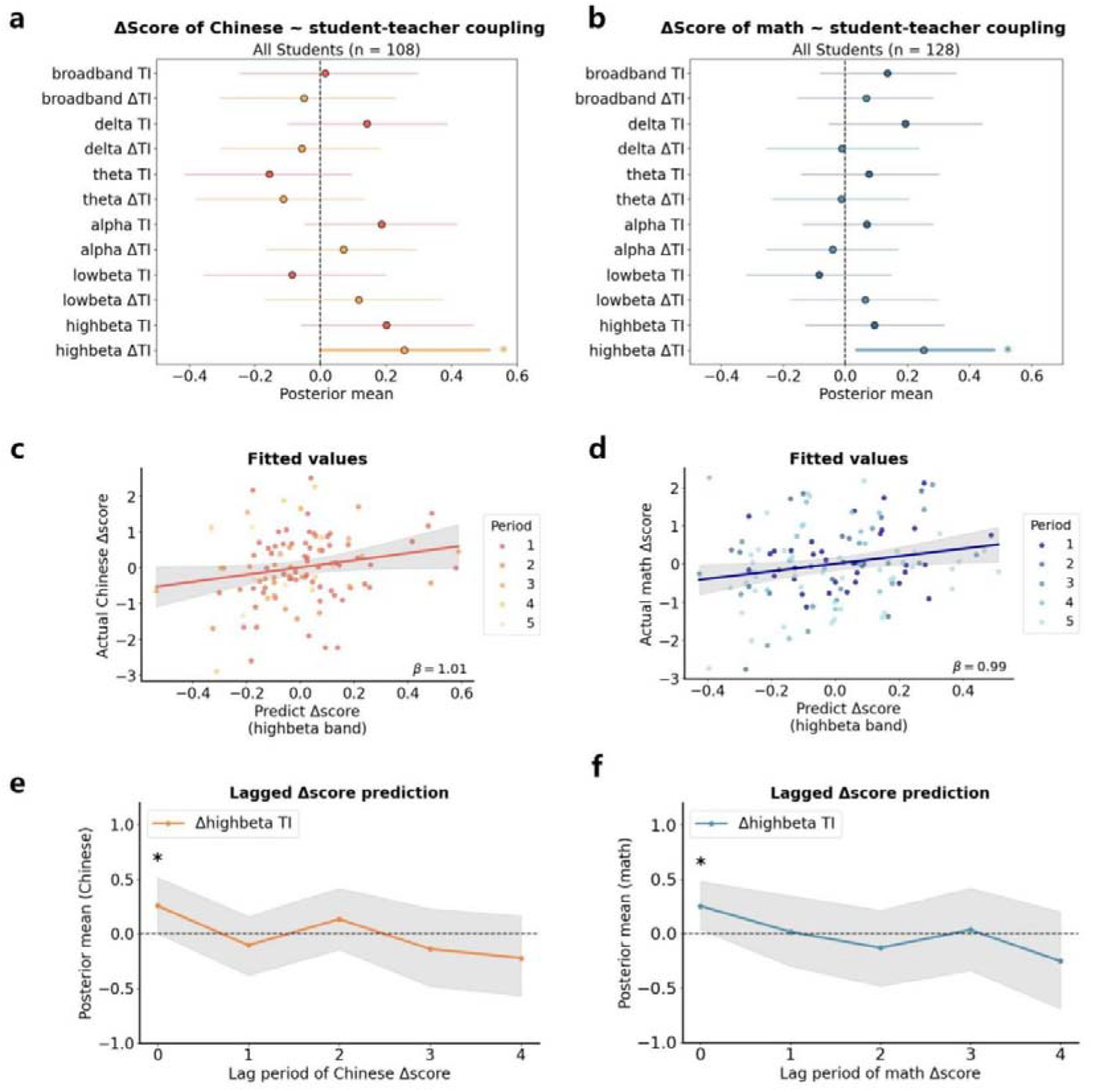
Bayesian estimation results of the dynamic model. **a**, Dynamic model estimates for Chinese (full sample, N = 108). The forest plots display the posterior mean estimates and their 95% highest posterior density intervals (HPDIs). The horizontal axis represents the posterior mean estimates, while the vertical axis indicates different frequency bands Asterisks denote significant correlation, *pl<l0.05, **p□<□0.01, ***p□<□0.001. **b**, Dynamic model estimates for math (full sample, N = 128). Visualization follows the same convention as in panel a. **c, F**itted values of forecasting Δscore of Chinese. The horizontal axis denotes predict Δscore, while the vertical axis represents the actual values of Δscores. Data points from different *periods* are denoted by distinct colors. **d**, Fitted values of forecasting Δscore of math. Axes and color-coding follow the same convention as in panel c. **e**, Lagged prediction of ΔChinese scores. The horizontal axis represents lag periods, and the vertical axis displays posterior mean estimates. Shaded areas indicate 95% HPDIs. **f**, Lagged prediction of Δmath scores. Formatting is consistent with panel e.

In the junior student subgroup, changes in both broadband and high-beta TI values significantly predicted improvements in Chinese scores (Δbroadband TI: posterior mean = 0.520, 95% HPDI [0.004, 1.063]; Δhigh-beta TI: posterior mean = 0.643, 95% HPDI [0.163, 1.114]), while only ΔTI in the high-beta band was significantly associated with math score gains (Δhigh-beta TI: posterior mean = 0.296, 95% HPDI [0.006, 0.584]). By contrast, no credible effects were observed in the senior student subgroup for either subject, as the 95% HPDIs included zero. TI values were also found to significantly predict academic improvement, especially within the junior student subgroup. For Chinese, TI values in the broadband and high-beta frequency bands similarly exhibit a significant positive predictive effect on score increases (broadband TI: posterior mean = 0.628, 95% HPDI [0.047, 1.142]; high-beta TI: posterior mean = 0.797, 95% HPDI [0.297, 1.276]). For math, TI values in the broadband and delta, and bands are capable of positively forecasting increases in scores (broadband TI: posterior mean = 0.369, 95% HPDI [0.077, 0.666]; delta TI: posterior mean = 0.389, 95% HPDI [0.056, 0.729]). This implies that higher coupling values with teachers can also account for the progress in learning outcomes. All statistically significant findings were consistently validated through both Bayesian and parametric estimation (see *Table S1, S2* and *Fig. S1* for full results).

To explore the lagged effect of student-teacher coupling on Δscores across multiple subsequent periods, TI and ΔTI were employed to forecast Δscores with lags ranging from 0 to 4 periods. For instance, if ΔTI represents the change in TI values from period 1 to period 2, then with a lag period of 0, the corresponding dependent variable would be the change in scores from period 1 to period 2, yielding results consistent with those depicted in Fig. 2a and 2b. When the lag period is set to 1, the dependent variable shifts to the change in scores from period 2 to period 3. The outcomes are depicted in Fig. 2e (Chinese) and Fig. 2f (math), with only the significant results at lag 0 displayed in the figures. The results suggests that student-teacher coupling does not have a predictive effect on subsequent changes in academic performance.

### Student-teacher inter-brain coupling forecast students’ academic achievement

As a comparison, the study implemented a *static model* to forecast the coefficients between learning outcomes within the same period and student-teacher inter-brain coupling (TI values). The results, as shown in Fig. 3a, indicate no significant association between inter-brain coupling and student performance in Chinese classes across different subsamples. However, in math classes, a significant negative association was observed between student-teacher inter-brain coupling and student performance (Fig. 3b). Specifically, in the full sample, theta-band TI was negatively associated with math scores (theta TI: posterior mean = −0.134, 95%, HPDI [-0.26, −0.013]). Among senior secondary students, significant negative associations were observed between math performance and TI in the delta, theta, and low-beta frequency bands (delta TI: posterior mean = −0.194, 95%, HPDI [-0.382, −0.007]; theta TI: posterior mean = −0.283, 95%, HPDI [-0.473, −0.088]; low-beta TI: posterior mean = −0.217, 95%, HPDI [-0.421, −0.008]), as shown in *Fig. S2*. Fig. 3c present the model fitting results for the frequency bands that showed significant effects in the static model. The model outcomes from both Bayesian and parametric estimations are presented in *Table S3, S4*. The study subsequently employed TI and ΔTI values to predict student performance with temporal lags ranging from 0 to 4 periods (Fig. 3d). No significant associations were identified when forecasting academic outcomes across multiple lagged periods.

**Fig. 3.**
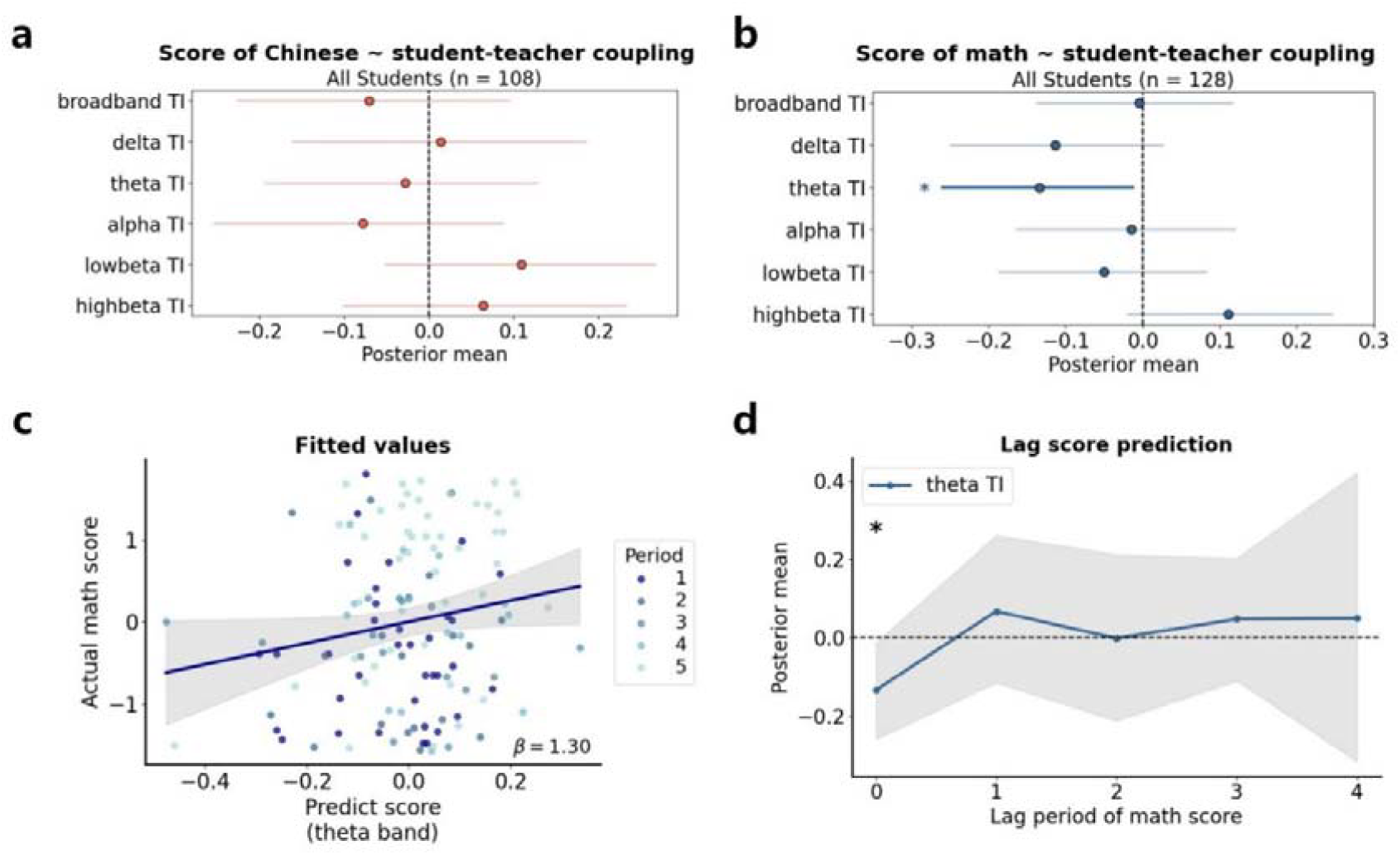
Bayesian estimation results of the static model. **a**, Static model estimates for Chinese (full sample, N = 108). Forest plots display the posterior mean estimates and their 95% HPDIs. The horizontal axis represents the posterior mean value of the estimatione, while the vertical axis indicates different frequency bands Asterisks denote significant correlation, *p□<□0.05, **p□<□0.01, ***p□<□0.001. **b**, Static model estimates for math (full sample, N = 128). **c**, The fitted values of forecasting math score. The horizontal axis shows predicted scores, and the vertical axis indicates actual scores. Data points from different time periods are distinguished by color. **d**, Lagged prediction of math scores. The horizontal axis represents lag periods, and the vertical axis displays posterior mean estimates. Shaded areas indicate 95% HPDIs.

In contrast to the dynamic model, the static model revealed significant negative associations between student–teacher inter-brain coupling and student performance within the same period. This pattern may reflect teachers’ adaptive instructional strategies, whereby increased attention is directed toward lower-performing students to help them keep pace with classroom instruction. However, from a longitudinal perspective, the dynamic model showed that students who experienced increases in coupling over time tended to exhibit corresponding improvements in academic achievement.

### Comparing the Predictive Effects of Student-Student and Student-Teacher Inter-brain Coupling on academic achievement

As active participants in classroom learning, students also engage in meaningful interactions with one another. Inter-brain coupling among peers has been shown to reflect shared attention and group-level engagement, with potential impacts on individual learning outcomes (Chen et al., 2023; Dikker et al., 2017). As a comparative analysis, we examined large-scale real-world EEG data to calculate the inter-brain coupling between individual students and their class-average peers, as well as between students and high-performing classmates. Both static and dynamic models were employed to assess the predictive value of these peer-based coupling measures for students’ academic outcomes.

The results of the dynamic model based on student-student coupling are presented in Fig. 4. Fig. 4a and 4b display the predictive effects of student-class inter-brain coupling on academic progress. No significant association was observed between student-class coupling and changes in Chinese scores. However, in the case of math, increases in alpha and low-beta Δ TI were positively associated with improvements in academic performance (alpha ΔTI: posterior mean = 0.208, 95%, HPDI [0.021, 0.377]; low-beta ΔTI: posterior mean = 0.217, 95%, HPDI [0.03, 0.412]). These results suggest that higher student-class inter-brain coupling in specific frequency bands may facilitate math learning in real-world classroom settings. Fig. 4c and 4d show the dynamic prediction results based on student-excellent inter-brain coupling. No significant associations were found between ΔTI and academic improvement across the full student samples. The analysis results for the sub-samples of junior and senior high school students are presented in *Fig. S3*, with trends consistent with those observed in the full sample analysis. Compared to student-teacher coupling, student-student inter-brain coupling showed a significant effect only in predicting improvements in math scores based on student-class coupling. In contrast, student-teacher coupling exhibited significant predictive power for academic growth in both Chinese and math subjects.

**Fig. 4.**
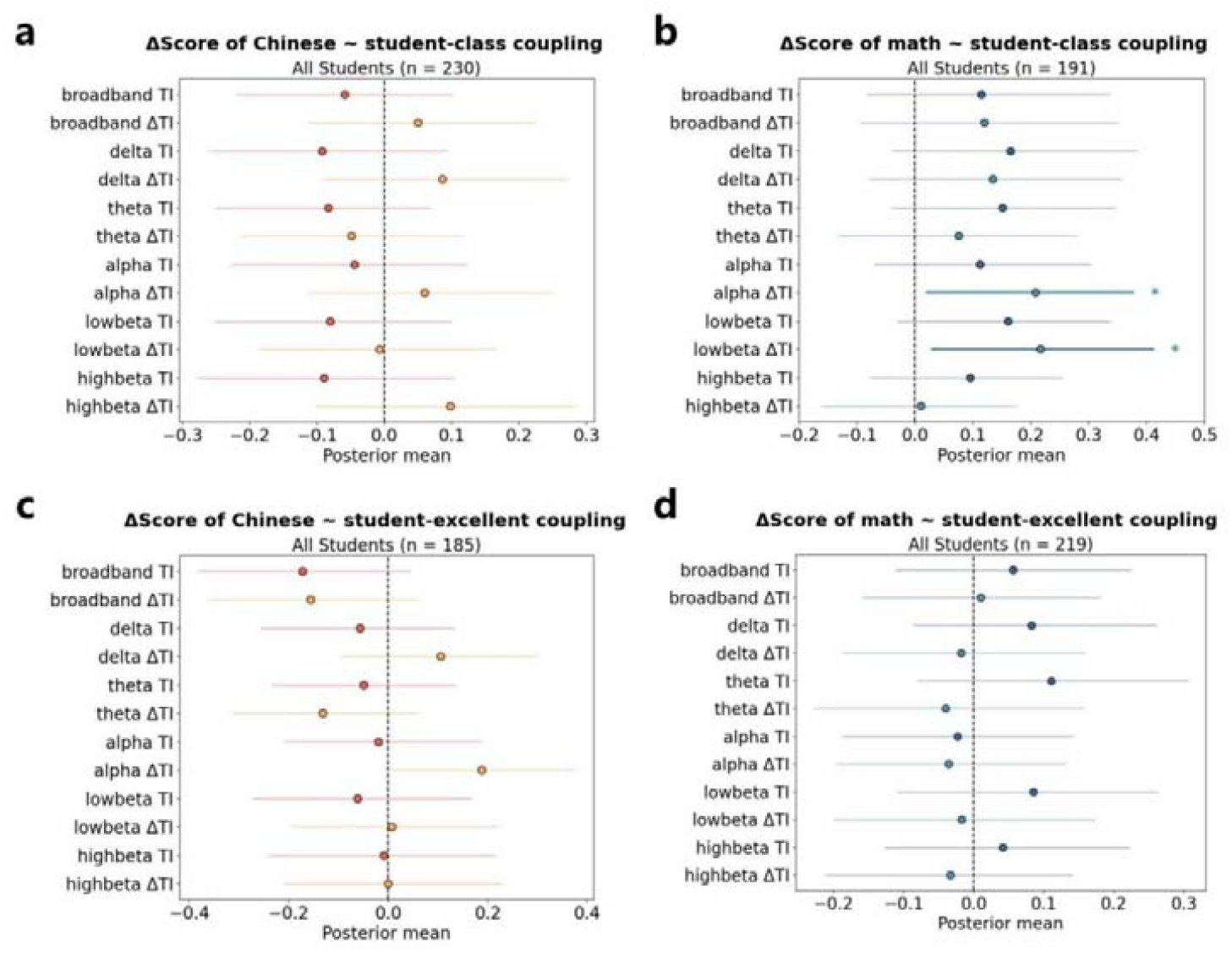
Bayesian estimation results of the dynamic model based on student-class coupling and student-excellent coupling. **a**, Dynamic model estimates for Chinese based on student-class inter-brain coupling (full sample, N = 230). **b**, Dynamic model estimates for math based on student-class inter-brain coupling (full sample, N = 191). **c**, Dynamic model estimates for Chinese based on student-excellent inter-brain coupling (full sample, N = 105). **d**, Dynamic model estimates for math based on student-excellent inter-brain coupling (full sample, N = 219).

The results of the static model based on student-class and student-excellent inter-brain coupling are presented in Fig. 5. As shown in Fig. 5a and 5b, no significant associations were observed between student-class coupling and Chinese performance. However, for math, student-class coupling in the low-beta frequency band significantly and positively predicted academic outcomes (low-beta TI: posterior mean = 0.095, 95%, HPDI [0.017, 0.176]). For student-excellent coupling, TI in the delta and alpha frequency bands was significantly positively associated with Chinese performance (delta TI: posterior mean = 0.107, 95%, HPDI [0.013, 0.199]; alpha TI: posterior mean = 0.113, 95%, HPDI [0.020, 0.199]), while alpha band TI was significantly negatively associated with math scores (alpha TI: posterior mean = −0.081, 95%, HPDI [-0.158, −0.007]). The analysis results for the sub-samples of junior and senior high school students are presented in *Fig. S4*, with trends consistent with those observed in the full sample analysis. Compared to the static model based on student-teacher coupling, both student-excellent and student-teacher inter-brain coupling exhibited similar negative associations with math performance, while student-class coupling revealed a distinct pattern of association. Additionally, the results demonstrated subject-specific patterns in neural coupling with high-performing peers, showing positive associations with Chinese performance but negative associations with math.

**Fig. 5.**
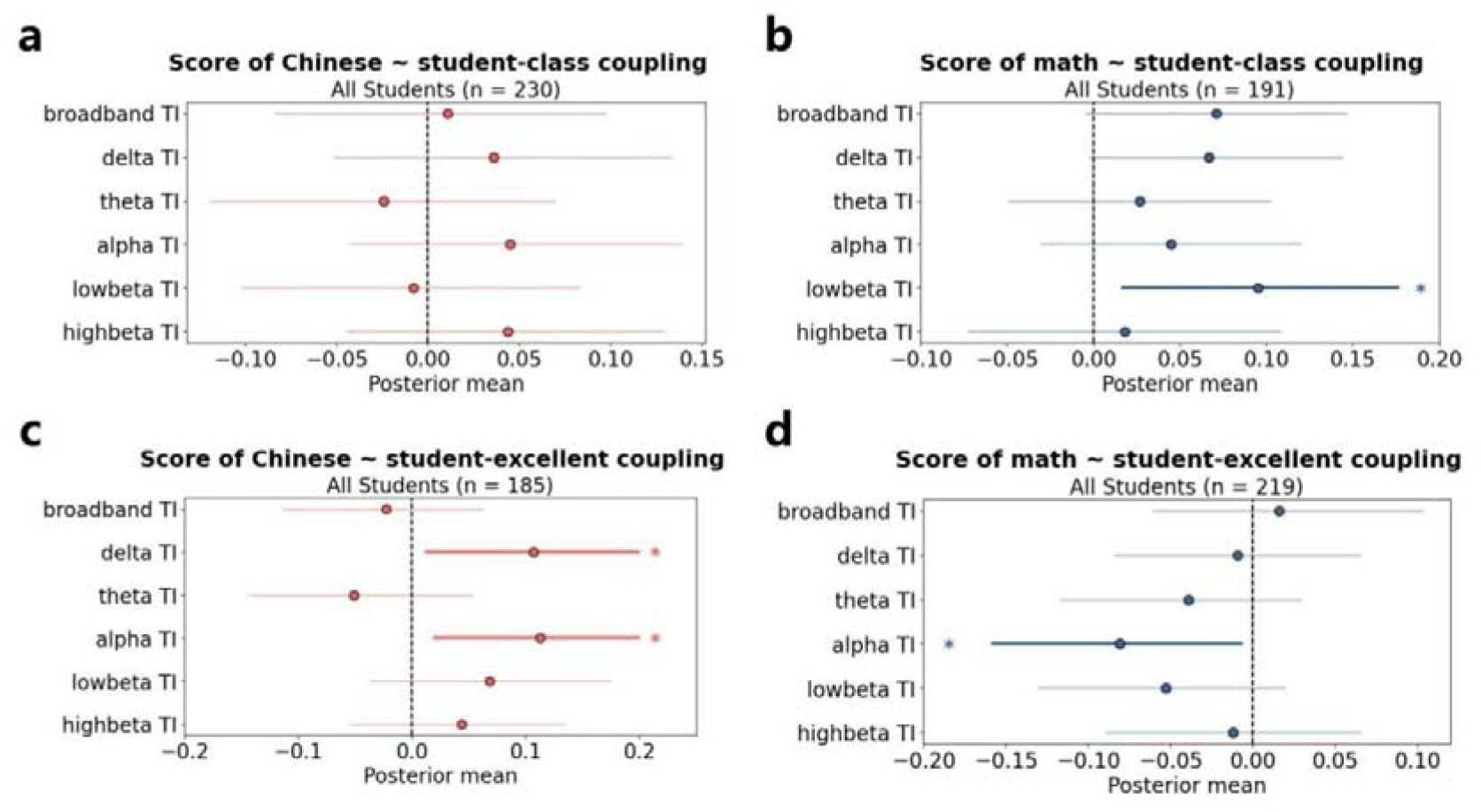
Bayesian estimation results of the static model based on student-class coupling and student-excellent coupling. **a**, Static model estimates for Chinese based on student-class inter-brain coupling (full sample, N = 230). **b**, Static model estimates for math based on student-class inter-brain coupling (full sample, N = 191). **c**, Static model estimates for Chinese based on student-excellent inter-brain coupling (full sample, N = 105). **d**, Static model estimates for math based on student-excellent inter-brain coupling (full sample, N = 219).

### Assessing the Subject Specificity of Student-Teacher Coupling Effects

To examine whether the predictive power of student-teacher inter-brain coupling is subject-specific, we conducted cross-subject analyses by using Chinese-class TI values to predict math performance and vice versa. Both dynamic and static models were applied. As shown in *Fig. S5*, no significant effects were found in either direction. This pattern suggests that the predictive effects of inter-brain coupling are not generalized across academic domains but instead reflect subject-specific dynamics of classroom interaction.

### Forecasting Learning Outcomes Across Different Student Performance Groups

Previous analyses revealed a negative association between student–teacher coupling and math achievement in the static model, suggesting that teachers may selectively attune more closely to lower-performing students during routine instruction. This suggests that the predictive relationship between neural coupling and academic performance may differ across achievement levels. To investigate this further, we categorized students into four groups based on their baseline performance: top 25% (75-100%), 50-75%, 25-50%, and bottom 25% (0-25%). Dynamic and static models were then estimated separately for each subgroup, and model fit indices were compared. As shown in Table 1, the predictive effects of student-teacher coupling varied across performance groups. For Chinese, the dynamic model yielded the most significant estimates within the top 25% of students. In contrast, for math, the best model fit in the dynamic analysis was observed among middle-performing students (25%-75%), while the static model showed the strongest fit for the bottom-performing group (bottom 25%). These findings confirm that the predictive association between student-teacher inter-brain coupling and academic performance differs across achievement levels, indicating that the relationship is modulated by students’ prior academic standing.

**Table 1.** The model fit for students at different performance levels.

|  | Chinese |  | Math |  |  |
| --- | --- | --- | --- | --- | --- |
|  | *Dynamic model* |  | *Dynamic model* |  | *Static model* |
|  | N | High-beta band<br>Wald chi2(2) | N | High-beta band<br>Wald chi2(2) | Theta band<br>Wald chi2(1) |
| ALL | 108 | 4.18 | 129 | 5.88 | **5.20** |
| 75%-100% | 25 | **4.03** | 32 | 4.95 | 0.08 |
| 25%-75% | 56 | 2.76 | 64 | **6.42** | 0.17 |
| 0%-25% | 27 | 1.00 | 33 | 3.95 | **0.30** |

## Discussion

This study investigates the impact of student-teacher inter-brain coupling values on learning outcome in authentic Chinese and math classes over six periods (i.e., three semesters) at a middle school. Using both static and dynamic models, we examined momentary associations and temporal fluctuations in student–teacher inter-brain coupling in relation to academic performance. The study found that increases in coupling over time significantly predicted academic improvement in both subjects, while static coupling values were negatively associated with math performance—likely reflecting teacher self-selection effects. No significant lagged effects were observed across models. Comparisons with student-student coupling revealed that student-class coupling predicted both short- and long-term math performance but showed no effect for Chinese, whereas coupling with high-performing peers was only associated with short-term outcomes. These findings underscore the critical role of student-teacher inter-brain coupling in shaping learning outcomes and offer empirical support for leveraging brain-based metrics to understand and optimize instructional interactions in real-world classrooms.

This study, through the application of dynamic modeling, reveals that students who experience enhanced inter-brain coupling with their teacher during class tend to achieve higher academic gains. Specifically, those students who benefit the most are those exhibiting elevated levels of neural coupling with their teachers, particularly when this coupling value shows significant enhancement. This finding underscores the practical significance of Δinter-brain coupling values as a valuable metric in real-world educational settings. By providing feedback to teachers and students regarding the changes in inter-brain coupling, this measure can offer timely, accurate insights into instructional efficacy, facilitating real-time adjustments. Fig. 6a and 6b illustrate the corresponding TI values and ΔTI values, respectively, when ranked by score and Δscore. Specifically, as shown in Fig. 6b, teachers can more intuitively observe that students who achieved greater academic improvements consistently exhibited higher increase in TI values.

**Fig 6.**
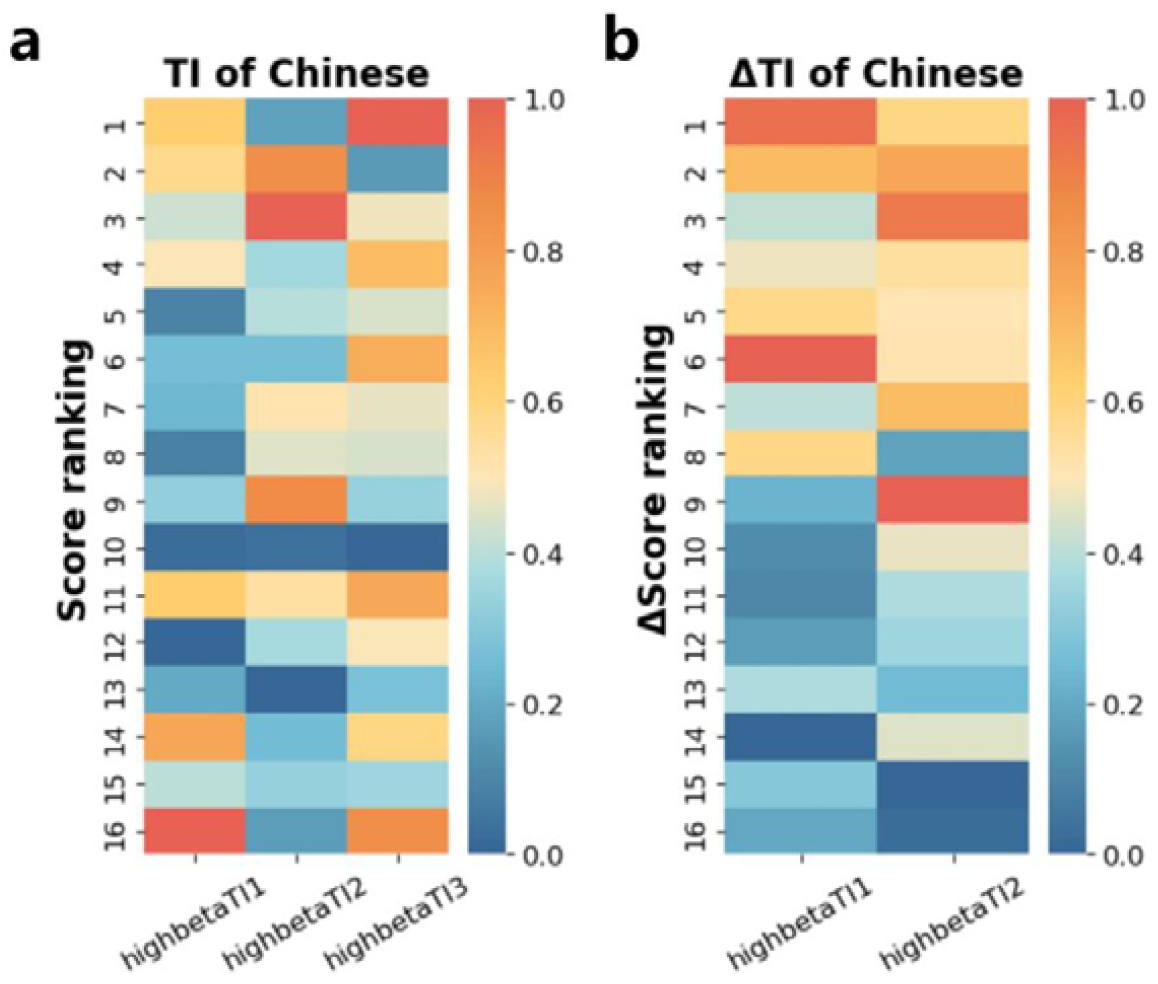
Visualization of student-teacher inter-brain coupling values across different score distributions. **a**, Heatmap of TI values sorted by Chinese score ranking. Students are arranged from top to bottom in descending order of their Chinese scores, with the x-axis corresponding to high-beta TI values across periods 1, 2, and 3. **b**, Heatmap of TI values sorted by predicted Chinese score. Students are arranged from top to bottom in descending order of their score improvements, with the x-axis corresponding to high-beta TI values across periods 1 and 2. Only students with data for all periods are included. Warmer colors represent higher values, while cooler colors represent lower values.

On the contrary, the static model, by identifying associations between student-teacher inter-brain coupling and concurrent academic performance, offers insight into short-term, surface-level relationships between coupling and learning outcomes. It reflects how teachers’ instructional strategies and attention distribution manifest in specific classroom contexts. In real classrooms, however, teachers often adopt adaptive interaction strategies based on students’ prior performance—a phenomenon known as teacher self-selection (Willis & Rosen, 1979). This tendency to engage more with lower-performing students to boost overall class outcomes may paradoxically result in higher coupling among students who perform worse academically, leading to negative associations in static models. This pattern was observed in our study, where student–teacher coupling in math classes was negatively correlated with achievement. Prior research similarly suggests that coupling values alone are insufficient for reliably forecasting academic outcomes (Bevilacqua et al., 2019), as static models may encounter uncontrolled confounding in predictive contexts. In contrast, the dynamic model, by capturing temporal fluctuations in coupling and linking them to longitudinal changes in academic outcomes, offers greater control over teacher-driven selection effects and student-level variability. This enables stronger causal inference about the benefits of high-quality student–teacher interaction.

To explore the unique advantage of student-teacher coupling, the study compared student-teacher inter-brain coupling with student-student coupling in predicting academic outcomes. Results showed that student – class coupling significantly predicted score fluctuations in math over time, suggesting a potential causal link. In contrast, student– excellence coupling was associated only with momentary performance and lacked longitudinal predictive power. Notably, these effects were subject-specific: student–class coupling was more predictive in math, while student–excellence coupling was more relevant in Chinese. These findings align with prior research suggesting that alignment with the class average supports learning in analytical subjects like math, whereas high-achieving peers may serve as better cognitive benchmarks in Chinese (Chen et al., 2023). Crucially, across both subjects, student-teacher inter-brain coupling consistently demonstrated the strongest and most stable predictive power—for both immediate performance and long-term improvement.

In addition to these findings, the main effects in the dynamic model were most prominent in the high-beta frequency band, whereas the static results were primarily associated with theta-band synchrony. This divergence suggests that different aspects of teacher-student interaction are captured by distinct neural dynamics. High-beta coupling has been linked to top-down cognitive control (Buschman & Miller, 2007; Kornblith et al., 2016), goal-directed processing (Karch et al., 2016), integration of complex information, and long-term memory (Hanslmayr et al., 2014), aligning with the notion that sustained improvement reflects deeper, effortful engagement (Baceviciute et al., 2021; Chikhi et al., 2022). In contrast, theta-band activity is associated with attentional allocation and episodic encoding (Guderian et al., 2009; Landau et al., 2015), reflecting more immediate, surface-level responsiveness to instruction. Thus, while high-beta coupling may signal long-term alignment in cognitive processes conducive to academic growth, theta synchrony may indicate moment-to-moment sensitivity to pedagogical input. These frequency-specific patterns underscore the multifaceted nature of classroom neural coupling and highlight the value of combining static, dynamic, and experimental approaches to capture the neural correlates of effective teaching. Notably, our findings revealed no significant lagged effects, suggesting that the predictive influence of inter-brain coupling is limited to contiguous academic terms—likely due to the dynamic nature of adolescent performance, with the impact of neural coupling most evident within the relevant teaching period.

Several limitations of this study warrant discussion. First, to minimize disruption to real classroom environments, we employed wearable, portable devices for data collection. However, these devices have limitations, particularly in the number of available channels. Future studies may benefit from using multimodal wearable equipment, integrating additional physiological signals such as skin conductance and heart rate to enhance model development and predictive accuracy. Second, variations in classroom structures—shaped by factors such as national, cultural, and grade-level differences—may also impact results, underscoring the need for more diverse measurements in future research.

In conclusion, this study systematically explored the impact of teacher-student coupling on learning outcomes in authentic classroom settings, providing neurophysiological evidence across multiple dimensions. Findings highlight the pivotal role of these interactions in enhancing learning and underscore the importance of increased student-teacher inter-brain coupling in predicting academic gains. For teachers, this increase offers an efficient way to assess instructional efficacy, while providing students with personalized feedback to understand their performance and adjust study strategies, demonstrating this approach’s practical value in education.

## Materials and Methods

### Participants

The study was a longitudinal observational study involving 141 students in their first year of junior and senior secondary school, along with eight teachers (four teaching Chinese and four teaching math) from four distinct classes at a middle school in Beijing. Of the original sample, 34 students withdrew during the study period. Ultimately, 107 students (60 males, 48 from junior secondary school, aged 12–15) who had valid and consecutive data across at least two time points were included in the final analyses.

All participants and their legal representatives provided informed consent, and the study was approved by the Department of Psychology at Tsinghua University (THU201708).

### Data Collection

#### EEG data

The EEG data collection took place in authentic classroom environment and spanned a total duration of 12 months, covering three academic semesters. Data acquisition extended over four months per semester, with each half-semester identified as a distinct ‘period’. Within these periods, data collection took place for a week every month, during which one to two sessions were recorded daily from four separate student classes. In total, 393 class sessions were recorded, including 182 Chinese classes (88 junior, 94 senior) and 211 math classes (113 junior, 98 senior). A total of 4,175 valid EEG recordings were obtained across sessions, including 2,057 from Chinese classes, with each student contributing one recording per session. Each lesson had a duration of 40 minutes, during which the teachers conducted their planned teaching activities, and students engaged in their regular learning tasks. Both students and teachers equipped the EEG headbands before each session and removed them subsequently, as directed by the research team.

EEG data were captured using a dual-channel headband featuring dry electrodes, specifically positioned at the Fp1 and Fp2 regions of the frontal cortex. The data recording occurred at a sampling rate of 250 Hz, employing the Brainno device (SOSO H&C, South Korea). The reference electrode was clipped to the right ear, serving as the ground for Fpz.

#### Student’s learning outcomes

Data on the students’ academic achievement were gathered through their midterm and final examination scores in Chinese and math over three semesters, encompassing a total of six examinations, provided by the school administration.

### Data Processing

#### Student’s learning outcome preprocessing

We utilized standardized ranking information to represent students’ learning outcomes, enabling a consistent and equitable comparison of student performance across various exams and cohort sizes. Students were first ranked on a scale from 1 to *N*, where *N* represents the total number of students in the cohort. The highest-performing student was assigned a rank of 1, progressing sequentially to the student with the lowest score, who received the rank of *N*. This approach effectively mitigating the issues of score saturation and truncation and highlights the relative performance among students, enhancing the robustness of the model. This initial ranking was subsequently standardized using a specific formula:

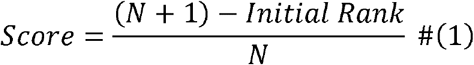

Where *score* denotes students’ standard score of students, representing their academic achievement. *N* denotes the total number of students, and the *Initial Rank* indicates the ranking of individual student within the cohort. The *score* ranges between 0 and 1, with higher values signifying better performance.

#### EEG data preprocessing

The EEG data for each student were preprocessed through a five-step procedure for both channels, consistent with methodologies employed in previous studies (Chen et al., 2023; Xu et al., 2024). The preprocessing steps included the identification of missing data, robust detrending (De Cheveigné & Arzounian, 2018), band-pass filtering, waveform segmentation, and ocular artifact mitigation. Subsequently, the data from each channel were segmented into 30-second epochs. For a single 40-minute class session, this resulted in a total of 160 epochs across both channels for every student (*Fig. S6*).

Additionally, 33 of the participants were also participates in a separate research project, during which eyes-open and eyes-closed EEG data in resting states were collected. The spectral analysis was conducted on resting state data. The spectrum of the eye-closed data prominently featured a peak in the alpha frequency band, thereby affirming the reliability of the collected EEG data (*Fig. S7*).

#### Student-teacher Inter-brain Coupling Calculation

This study employs the total interdependence values (TI) for assessing student-teacher, student-class and student-excellence inter-brain coupling. This approach relies on the coherence of two signals within the frequency domain, which has been validated for its efficacy (Xu et al., 2024). The utility of TI values as a metric for inter-brain coupling has been increasingly recognized in a variety of real-world contexts, including concert hall, and classroom (Bevilacqua et al., 2019; Chabin et al., 2022; Chen et al., 2023; Dikker et al., 2017). The computation of TI values is as follow:

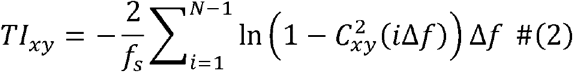

where *C*_*xy*_ denotes the spectral coherence of the signal at the frequency *f*, computed by the Welch method. Here, *f*_*s*_ denotes the sampling rate, and *N* represents the number of data points between 0 and the Nyquist sampling rate *f*_*s*_ /2. The frequency resolution Δ*f* is computed employing the relation 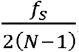. In this analysis, TI values were computed across the EEG’s broad frequency spectrum, ranging from 1 to 40Hz. Calculations also incorporated individual frequency bands: delta (1-4Hz), theta (4-8Hz), alpha (8-13Hz), low beta (13-18Hz) and high-beta (18-30Hz).

Within each session, the TI value for each student-teacher pair was determined by averaging the TI values across all epochs that successfully matched between the student and teacher, considering data from both recording electrodes (as shown in Fig. 1C). A ‘segment’ was defined as a matched epoch, recognized as such only when it included artifact-free and valid data from both the teacher and the student. A minimum threshold was established, requiring that only student-teacher pairs with at least six valid segments were to be included (Chen et al., 2023; Dikker et al., 2017). For Chinese classes, the average valid student-teacher coupling proportion per session was 47.2%, compared to 49.1% for math classes (*Fig. S8a, b*).

#### Student-class inter-brain coupling

The student-class inter-brain coupling value for a given student is computed as the mean of all TI values between this student and the rest of the class of each session. For Chinese, the average proportion of valid student-class coupling segments for each session was 57.8%, while for math, it was 57.1% (*Fig. S8c, d*).

#### Student-excellence inter-brain coupling

The student-excellence inter-brain coupling value for a given student is computed as the mean of all TI values between this student and the top four student in the class. For Chinese, the average proportion of valid student-class coupling segments for each session was 37.5%, while for math, it was 45.9% (*Fig. S8e, f*).

Following this, the inter-brain coupling values within each period (i.e., half-semester) were averaged for each student. To ensure the quality of the data, student samples with valid sessions constituting less than 20% within a period were omitted from the analysis. These averaged TI values were then normalized to a range of [0, 1] for each frequency band.

#### Regression Analysis

Linear mixed-effects (LME) models were fitted to the data to investigate the relationship between student-teacher TI and academic achievement, which allow to effectively partition the overall variation of the dependent variable into components corresponding to different levels of data hierarchy (Gałecki et al., 2013). According to the dependent variables, two kinds of models are designed. The *dynamic model* (formula 4) uses the difference in test score between *period* t and the previous *period* as the dependent variable, representing the improvement of learning achievement, while the *static model* (formula 5) uses standardized test score as dependent variable. Given the potential non-independence of EEG data within each individual students or time *period*, the two categorical variables are employed to classify the observations into separate groups, to account for correlated observations in the data (Pinheiro & Bates, 2000).

Bayesian estimation was employed to analyze the relationship between student–teacher inter-brain coupling and academic performance, using Markov Chain Monte Carlo (MCMC) sampling to approximate the posterior distributions of model parameters (Howson & Urbach, 2006; Humphreys & Jacobs, 2015). This approach was chosen for its flexibility in estimating complex hierarchical models and its ability to provide full posterior distributions of parameters, offering richer information than traditional point estimates. Bayesian methods do not rely on large-sample asymptotics or strict distributional assumptions, making them particularly suited for small to moderate sample sizes and nested data structures frequently encountered in educational neuroscience research.

To ensure the robustness of our reported effects, we adopted a dual reporting strategy: all significant findings reported in this paper meet the threshold of both Bayesian credible intervals (95% HDIs excluding zero) and frequentist nonparametric p-values (p < 0.05). This convergence across inferential paradigms enhances the credibility and interpretability of our results.

#### The intraclass correlation analysis

The intraclass correlation coefficient (ICC) was first calculated for each categorical variable, in order to determine if it is appropriate to include the corresponding random effect. The formula of ICC is as follows:

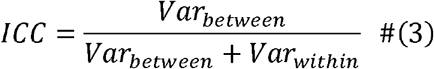

where *Var*_*between*_ denotes the between-group variation of the dependent variable, and *Var*_*within*_ denotes the within-group variation. The ICC value is between 0 and 1, where a higher ICC value indicates larger distinctions between groups. An ICC value above 0.05 highlights noteworthy intraclass correlation that should be considered (Bujang & Baharum, 2017).

#### Forecasting of academic improvement

The *dynamic* model is applied in order to investigate whether student-teacher coupling can forecast student academic achievement. The investigation aimed to explore whether the increase in TI values could dynamically affect changes in academic achievement after controlling for the initial impact of TI. In pursuit of this, the model integrated both the baseline TI values and their incremental growth as independent variables, with the growth of standardized scores as the dependent variable in the analysis.

The *dynamic models* are:

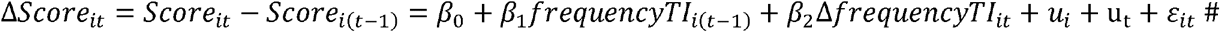

where Δ*score*_*it*_ denotes the improvement of students’ academic achievement; *score*_*it*_ represents the standardized score of student *i* in period *t*, and Score_i (t − 1)_ denotes the standardized score of the same student *i* in the preceding period, *t*−1; *frequencyTI*_st_*i*(*t* −1)_ indicate the vector of student-teacher TI values of student *i* in period *t*−1 in different frequency bands (i.e., broadband, delta, theta, alpha, low-beta and high-beta bands); Δ *frequencyTI*_*it*_ is the difference between *frequencyTI*_*it*_ and *frequencyTI*_*i,t* −1_; *u*_*i*_ is the individual fixed ability (or say, random effect of student), *u*_*t*_ is the period fixed characteristics (or say, random effect by period), and *ε*_*it*_ denotes the random error term.

ICCs were first calculated using the full student sample. For Δscore of Chinese, both the ICC for student individual (*ICC*_*stu_id*_) and for period factor (*ICC*_*period*_) was 0.000, indicating negligible variance attributable to these factors. For Δscore of math, *ICC*_*stu_id*_ was also 0.000, while *ICC*_*period*_ was 0.007, suggesting a small proportion of variance associated with temporal differences. In the subsample analysis, both *ICC*_*stu_id*_ and *ICC*_*period*_ for Chinese remained 0.000 in junior and senior secondary students. For math, *ICC*_*stu_id*_ was 0.137 in the junior group and 0.007 in the senior group, while *ICC*_*period*_ remained 0 in both groups. ICCs above 0.05 were treated as evidence of between-group variance and were included in the analytical model.

#### Forecasting of student’s academic achievement

The study then investigates the influence of student-teacher inter-brain coupling on the students’ learning outcome. The inter-class correlation analysis (ICC) was first conducted to refine the regression models. For Chinese score, the ICC for individual student (*ICC*_*stu_id*_) was 0.648, and for *period* (*ICC*_*period*_) it was 0.016; For math score, *ICC*_*stu_id*_ was 0.747, and *ICC*_*period*_ was 0.134. ICC values exceeding 0.05 were treated as random effects in the analysis.

The *static model* was designed as:

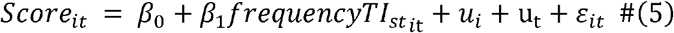

where all the symbols are consistent with those defined before (*See formula 4*).

#### Forecasting of student’s academic achievement over multiple lagged periods

To investigate whether the student-teacher coupling has a long-term predictive effect on score and Δscore, this study utilized current TI and ΔTI to forecast scores over multiple lagged periods. The formula is as follow:

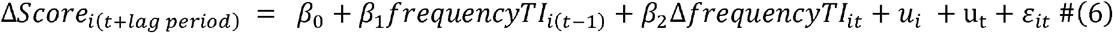

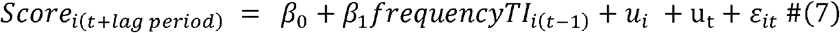

where *lag period* denotes the number of periods by which the score lags behind ΔTI, *score*_*i* (*t* + *lag period*)_ is the achievement of student i in period *t* + *lag period*; and other symbols are consistent with before (*See formula 4, 5*). For example, if the *lag period* is set to 2, and *t* equals 3, this implies forecasting the score for the 5th period using the TI from the 2nd period and the ΔTI from the 2nd to 3rd period.

## Acknowledgements and funding sources

This work was supported by the National Natural Science Foundation of China (62177030, T2341003, 61977041) and the Beijing Educational Science Foundation of the Fourteenth 5-year Planning (BGEA23019). Appreciation is extended to Xinqiao Gao, Baosong Li, Fei Qin, Xuan Qi, Yingying Zhao, Zhilin Qu, Tingting Wang, Xinya Liu and Kun Wang for their support during the data collection.

## Author Contributions

Xiaomeng Xu was responsible for the study’s conceptual design, methodology, data analysis, and writing the original manuscript. Dan Zhang contributed to the validation of the research and participated in the manuscript’s review and editing process. Yu Zhang led the research design, validated the findings, and engaged in reviewing and editing the manuscript.

## Additional Information

Supplementary Information is available for this paper. Correspondence and requests for materials should be addressed to Yu Zhang and Dan Zhang.

## Data Availability and Code Availability statements

The datasets generated during and/or analysed during the current study are not publicly available because the data are confidential but are available from the corresponding author on reasonable request.

