## Supplemental Information for "Syncing Minds for Learning: Student-Teacher Inter-brain Coupling Forecast Cross-semester Academic Fluctuation in Real-World Classrooms"

### Extended Data Figure

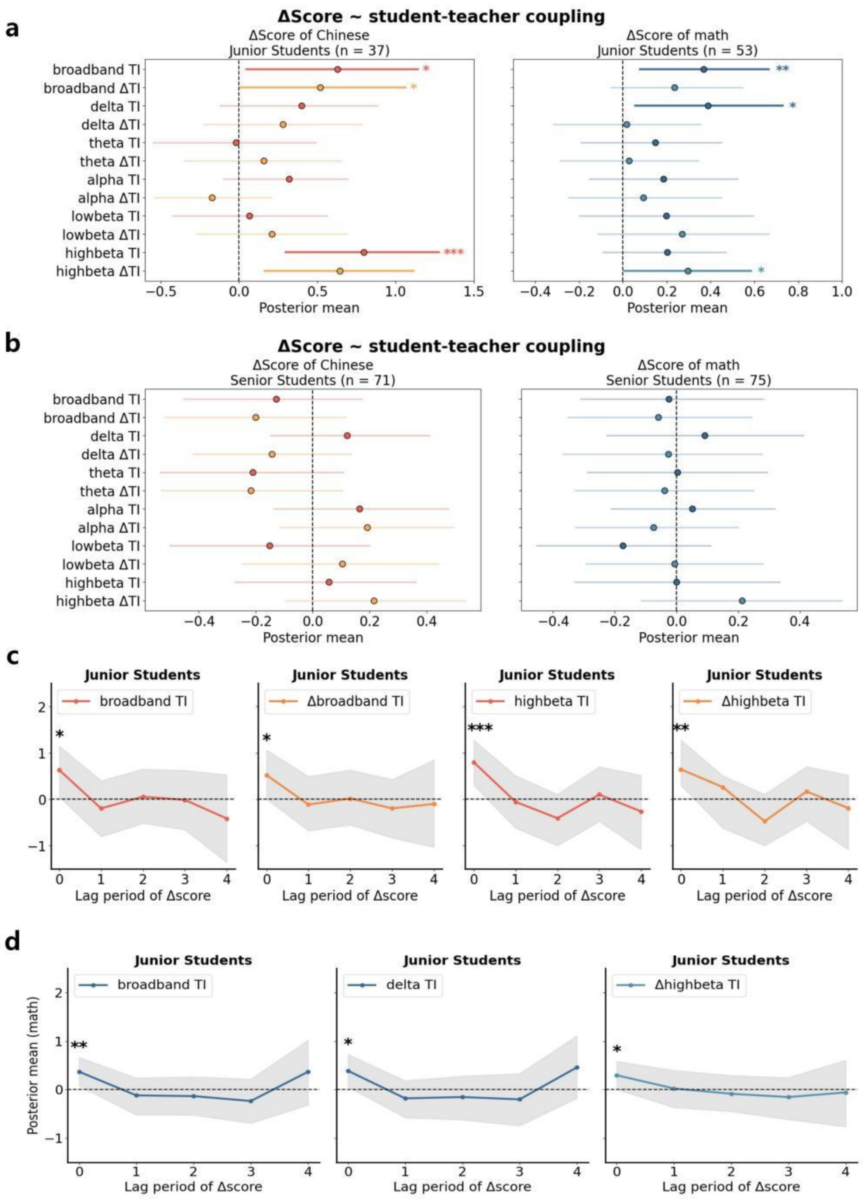


**Fig. S1. Bayesian estimation results of the dynamic model based on junior and senior high school student samples. a,** Dynamic model estimates for the junior high school student samples. The left panel shows the results for Chinese, and the right panel shows the results for math. The forest plots display the posterior mean estimates and their 95% highest posterior density intervals (HPDIs). The horizontal axis represents the posterior mean estimates, while the vertical axis indicates different frequency bands. Asterisks denote significant correlations: *$p < 0.05$, **$p < 0.01$, ***$p < 0.001$. **b,** Dynamic model estimates for the senior high school student samples. The left panel shows the results for Chinese, and the right panel shows the results for math. Visualization follows the same convention as in panel a. **c,** Lagged prediction of ΔChinese scores. The horizontal axis represents lag periods, and the vertical axis displays the posterior mean estimates. Shaded areas indicate the 95% HPDIs. **d,** Lagged prediction of Δmath scores. The visualization follows the same structure as in panel c, showing lagged predictions for the math sample.

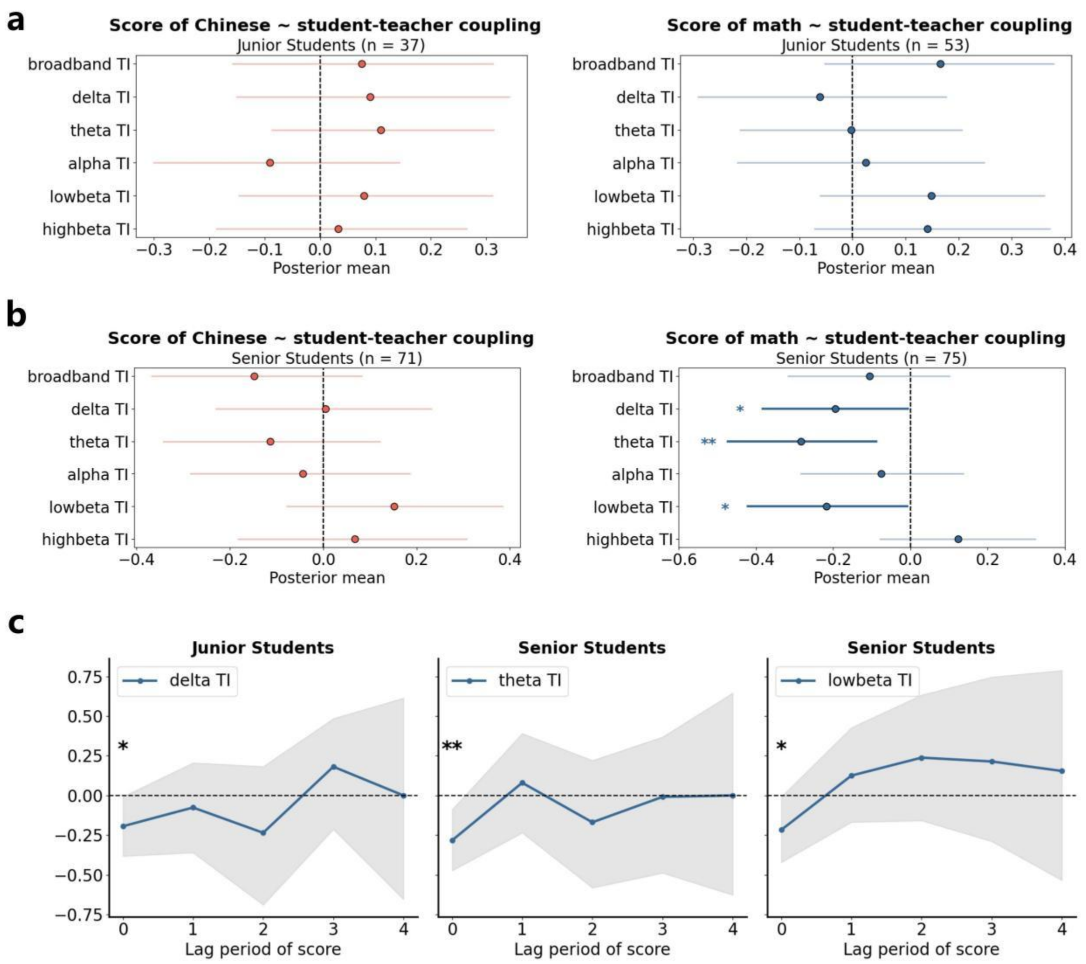


**Fig. S2. Bayesian estimation results of the static model based on junior and senior high school student samples. a,** Static model estimates for the junior high school studnets sample. The left panel shows the results for Chinese, and the right panel shows the results for math. The forest plots display the posterior mean estimates and their 95% highest posterior density intervals (HPDIs). The horizontal axis represents the posterior mean estimates, while the vertical axis indicates different frequency bands. Asterisks denote significant correlations: *$p < 0.05$, **$p < 0.01$, ***$p < 0.001$. **b,** Static model estimates for the senior high school student samples. The left panel shows the results for Chinese, and the right panel shows the results for math. Visualization follows the same convention as in panel a. **c,** Lagged prediction of math scores. The horizontal axis represents lag periods, and the vertical axis displays the posterior mean estimates. Shaded areas indicate the 95% HPDIs.

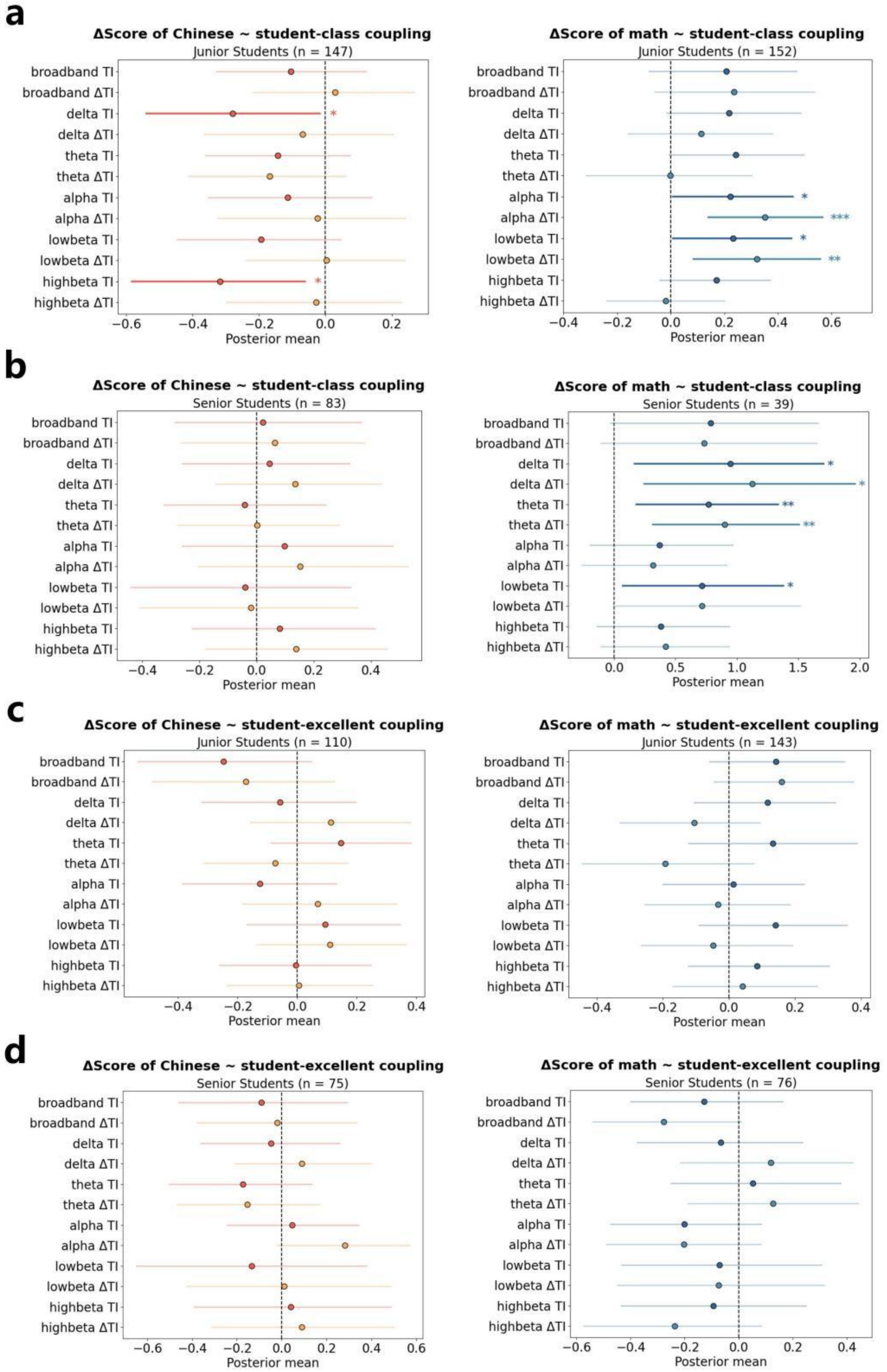


**Fig. S3. Bayesian estimation results of the dynamic model based on student-class coupling and student-excellence coupling, derived from junior and senior high school student samples.** **a,** Dynamic model estimates for Chinese (left) and math (right) based on student-class inter-brain coupling, derived from junior school student samples. **b,** Dynamic model estimates for Chinese (left) and math (right) based on student-class inter-brain coupling, derived from senior school student samples. **c,** Dynamic model estimates for Chinese (left) and math (right) based on student-excellent inter-brain coupling, derived from junior school student samples. **d,** Dynamic model estimates for Chinese (left) and math (right) based on student-excellent inter-brain coupling, derived from senior school student samples.

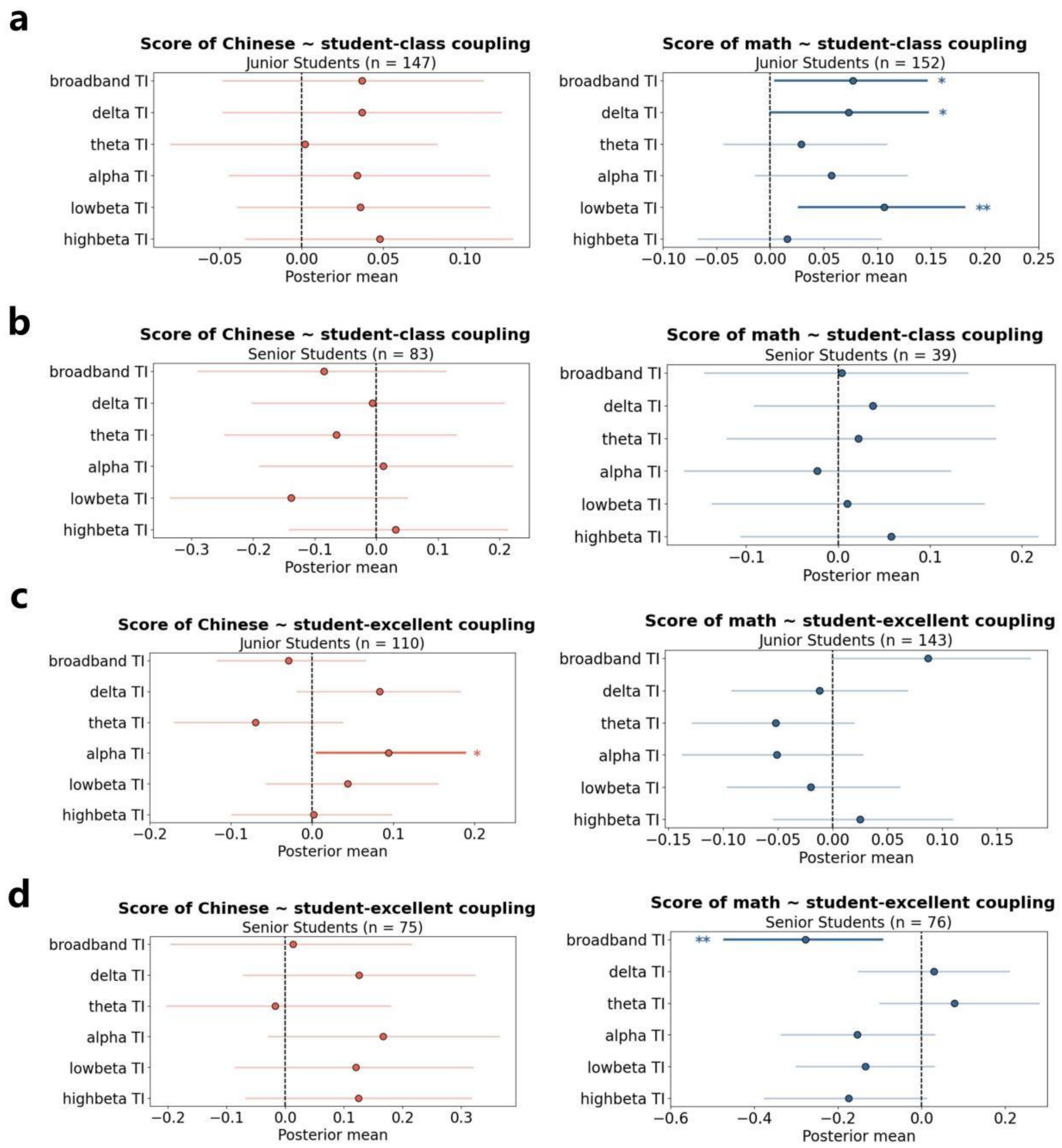


**Fig. S4. Bayesian estimation results of the static model based on student-class coupling and student-excellence coupling, derived from junior and senior high school student samples. a,** Static model estimates for Chinese (left) and math (right) based on student-class inter-brain coupling, derived from junior school student samples. **b,** Static model estimates for Chinese (left) and math (right) based on student-class inter-brain coupling, derived from senior school student samples. **c,** Static model estimates for Chinese (left) and math (right) based on student-excellent inter-brain coupling, derived from junior school student samples. **d,** Static model estimates for Chinese (left) and math (right) based on student-excellent inter-brain coupling, derived from senior school student samples.

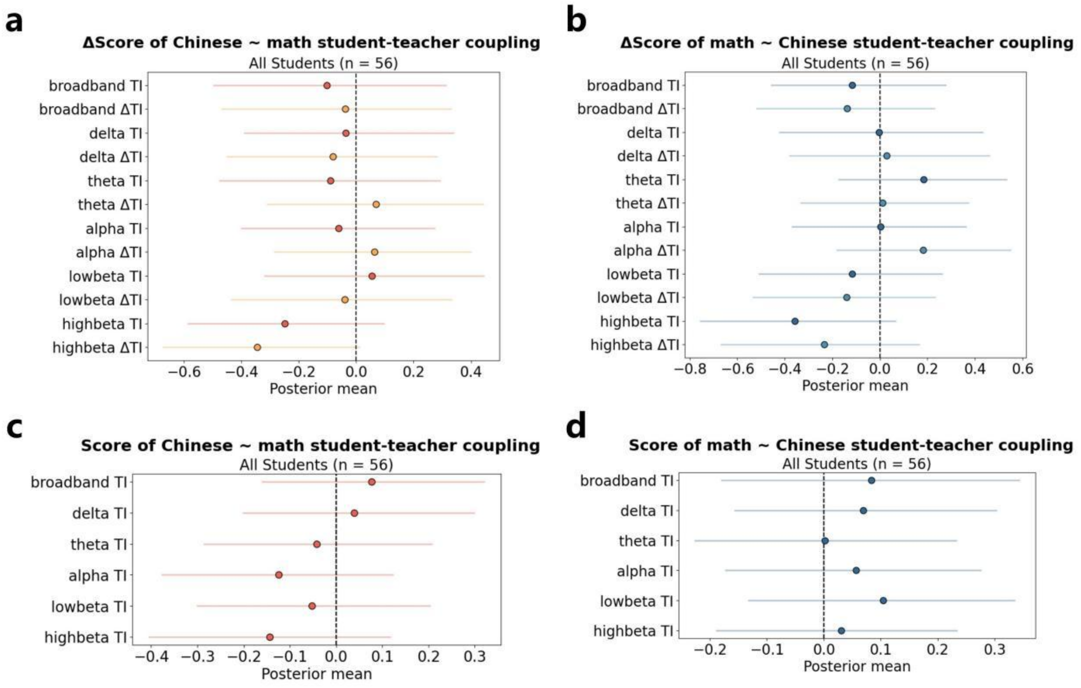


**Fig. S5. Cross-subject analyses of both dynamic and static models. a,** Dynamic prediction of Δ Chinese scores based on student-teacher coupling in the math class. **b,** Dynamic prediction of Δ math scores based on student-teacher coupling in the Chinese class. **c,** Static prediction of Chinese scores based on student-teacher coupling in the math class. **d,** Static prediction of math scores based on student-teacher coupling in the Chinese class.

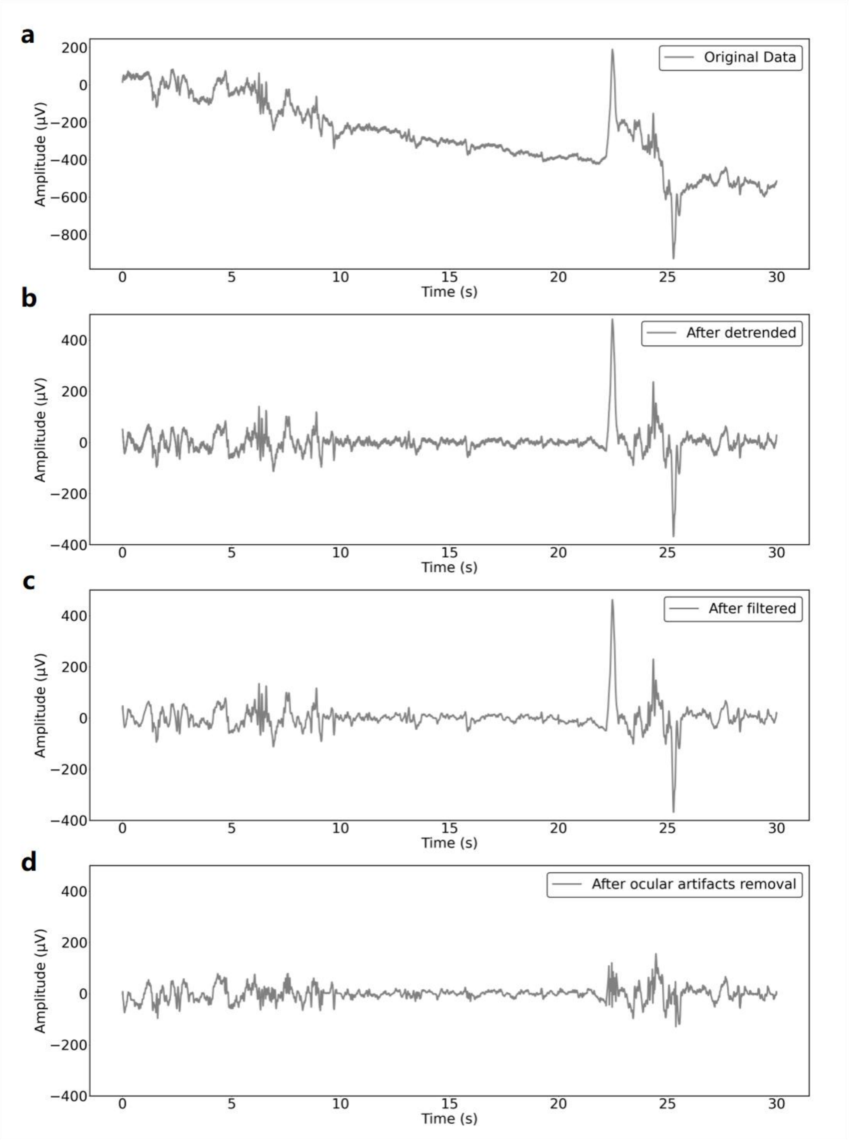


**Fig. S6.** An illustration of a representative EEG epoch preprocessing. For each student, EEG data from both the left and right channels, recorded over a 40-minute class session, underwent individual preprocessing. The data from each channel were separately segmented into 80 epochs, resulting in a total of 160 epochs per student, with each epoch lasting 30 seconds. During preprocessing, continuous signals exhibiting amplitudes below a threshold of 0.1 were categorized as missing data. Any epochs containing more than 50% missing data were considered unsuitable for analysis and were consequently discarded. A polynomial fitting method was then employed for detrending the data, encompassing the identification and marking of artifact-contaminated points. Artifacts and electrooculograms were then detected and carefully removed by the MSDL toolbox. Following this, the EEG data were subjected to a band-pass filter ranging from 1 to 40 Hz. In the final phase of preprocessing, any EEG epochs that exhibited peak amplitudes beyond the threshold of $\pm 150\ \mu V$ were excluded from further analysis, ensuring the retention of only high-quality data. **a,** An illustration of a representative EEG epoch of original data. **b,** An illustration of a representative EEG epoch after the detrending. **c,** An illustration of a representative EEG epoch after the filtering. **d,** An illustration of a representative EEG epoch after the ocular artifact removal procedure.

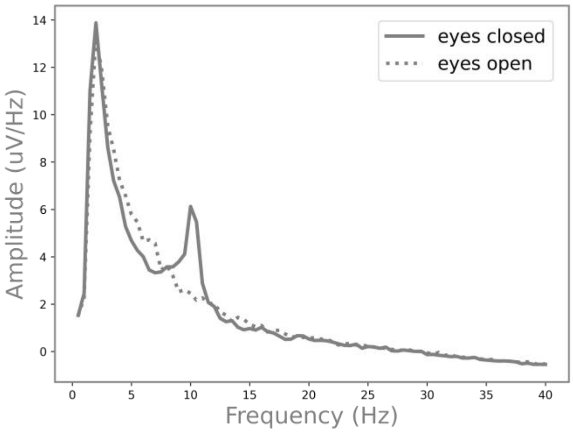


**Fig. S7.** Spectral analysis of eye-open/closed resting state EEG data from representative samples. The spectrum of the eye-closed data prominently featured a peak in the alpha frequency band, thereby affirming the reliability of the collected EEG data.

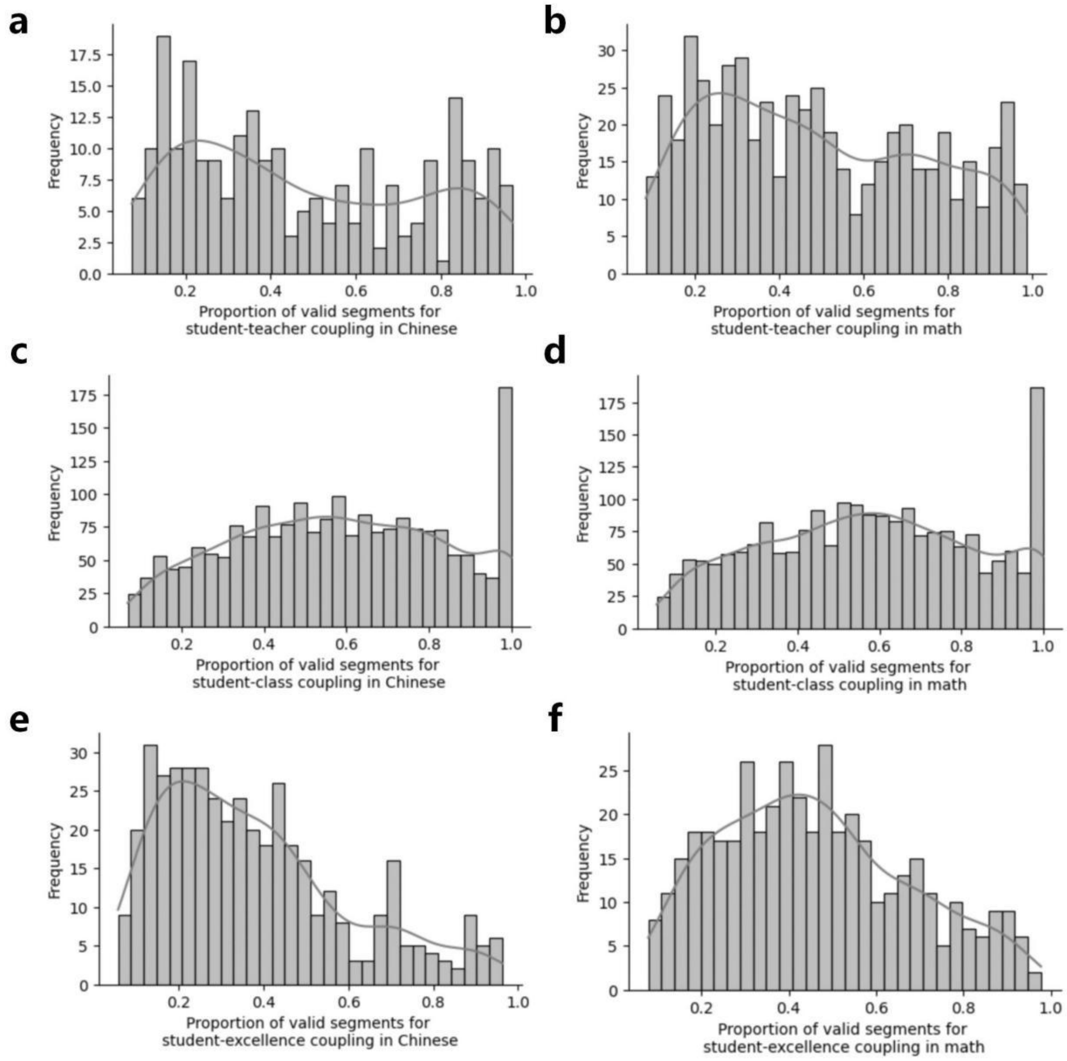


**Fig. S8.** Distribution of the proportion of valid coupling segments. **a,** Distribution of the proportion of valid student-teacher coupling segments to the number of valid EEG epochs of teacher in Chinese class. **b,** Distribution of the proportion of valid student-teacher coupling segments to the number of valid EEG epochs of teacher in math class. **c,** Distribution of the proportion of valid student-class coupling segments to the max number of valid EEG epochs of students in Chinese class. **d,** Distribution of the proportion of valid student-class coupling segments to the max number of valid EEG epochs of students in math class. **e,** Distribution of the proportion of valid student-excellent coupling segments to the max number of valid EEG epochs of top four students in Chinese class. **f,** Distribution of the proportion of valid student- excellent coupling segments to the max number of valid EEG epochs of top four students in math class.

**Table S1.** Parametric estimation of dynamic model results for Δscore based on student-teacher coupling. For the full sample, N = 108 for Chinese and N = 128 for math. For the junior high school student sample, N = 37 for Chinese and N = 53 for math. For the senior high school student sample, N = 71 for Chinese and N = 75 for math.

| | | **Δ Chinese Score** | **Wald chi2 (2)** | **Δ Math Score** | **Wald chi2 (2)** |
|---|---|---|---|---|---|
| ***All student sample*** | | | | | |
| **Broadband** | TI | 0.022<br>(0.145) | 0.36 | 0.142<br>(0.116) | 1.59 |
| | ΔTI | -0.043<br>(0.144) | | 0.080<br>(0.126) | |
| **Delta** | TI | 0.204<br>(0.166) | 3.58 | 0.265<br>(0.173) | 5.06 |
| | ΔTI | -0.058<br>(0.133) | | -0.018<br>(0.160) | |
| **Theta** | TI | -0.176<br>(0.148) | 1.44 | 0.086<br>(0.151) | 0.86 |
| | ΔTI | -0.134<br>(0.149) | | -0.024<br>(0.123) | |
| **Alpha** | TI | 0.221<br>(0.135) | 2.82 | 0.082<br>(0.111) | 1.29 |
| | ΔTI | 0.078<br>(0.129) | | -0.044<br>(0.127) | |
| **Low-beta** | TI | -0.084<br>(0.141) | 3.71 | -0.122<br>(0.156) | 2.29 |
| | ΔTI | 0.129<br>(0.154) | | 0.063<br>(0.144) | |
| **High-beta** | TI | 0.247<br>(0.149) | **4.18** | 0.104<br>(0.133) | **5.97** |
| | ΔTI | **0.297***<br>**(0.147)** | | 0.231<br>(0.101) | |
| ***Junior high scholl student sample*** | | | | | |
| **Broadband** | TI | **0.516***<br>**(0.202)** | **6.68** | **0.366****<br>**(0.143)** | **6.89** |
| | ΔTI | **0.403***<br>**(0.180)** | | 0.272<br>(0.168) | |
| **Delta** | TI | 0.416<br>(0.253) | 2.95 | **0.367***<br>**(0.156)** | **10.73** |
| | ΔTI | 0.235<br>(0.200) | | 0.012<br>(0.153) | |
| **Theta** | TI | 0.132<br>(0.211) | 0.83 | 0.140<br>(0.146) | 1.31 |
| | ΔTI | 0.291<br>(0.221) | | 0.026<br>(0.161) | |

| | | | | | |
|---|---|---|---|---|---|
| **Alpha** | TI | 0.263<br>(0.157) | 8.06 | 0.155<br>(0.145) | 1.27 |
| | ΔTI | -0.176<br>(0.118) | | 0.075<br>(0.156) | |
| **Low-beta** | TI | 0.065<br>(0.176) | 1.07 | 0.258<br>(0.235) | 2.44 |
| | ΔTI | 0.164<br>(0.178) | | 0.336<br>(0.223) | |
| **High-beta** | TI | **0.758*****<br>**(0.209)** | **13.17** | 0.181<br>(0.123) | **4.72** |
| | ΔTI | **0.504****<br>**(0.171)** | | **0.237***<br>**(0.114)** | |
| ***Senior high scholl student sample*** | | | | | |
| **Broadband** | TI | -0.157<br>(0.199) | 1.56 | 0.001<br>(0.170) | 0.11 |
| | ΔTI | -0.243<br>(0.199) | | -0.041<br>(0.150) | |
| **Delta** | TI | 0.213<br>(0.240) | 3.74 | 0.100<br>(0.250) | 0.79 |
| | ΔTI | -0.173<br>(0.173) | | -0.052<br>(0.212) | |
| **Theta** | TI | -0.284<br>(0.215) | 2.36 | -0.003<br>(0.226) | 0.19 |
| | ΔTI | -0.301<br>(0.214) | | -0.087<br>(0.187) | |
| **Alpha** | TI | 0.222<br>(0.197) | 1.89 | 0.069<br>(0.154) | 0.94 |
| | ΔTI | 0.244<br>(0.193) | | -0.089<br>(0.151) | |
| **Low-beta** | TI | -0.174<br>(0.206) | 3.84 | -0.205<br>(0.161) | 2.19 |
| | ΔTI | 0.135<br>(0.242) | | -0.037<br>(0.202) | |
| **High-beta** | TI | 0.084<br>(0.217) | 2.38 | 0.020<br>(0.212) | 3.47 |
| | ΔTI | 0.300<br>(0.217) | | 0.205<br>(0.150) | |

**Table S2.** Bayesian estimation of dynamic model results for Δscore based on student-teacher coupling. For the full sample, N = 108 for Chinese and N = 128 for math. For the junior high school student sample, N = 37 for Chinese and N = 53 for math. For the senior high school student sample, N = 71 for Chinese and N = 75 for math.

| | | **Δ Chinese Score** | | **Δ Math Score** | |
|---|---|---|---|---|---|
| | | **Posterior mean** | **HPDI 95%** | **Posterior mean** | **HPDI 95%** |
| ***All student sample*** | | | | | |
| **Broadband** | TI | 0.014 (0.138) | [-0.245, 0.295] | 0.135 (0.112) | [-0.079, 0.355] |
| | ΔTI | -0.051 (0.137) | [-0.305, 0.228] | 0.068 (0.111) | [-0.153, 0.28] |
| **Delta** | TI | 0.141 (0.124) | [-0.098, 0.385] | 0.194 (0.124) | [-0.051, 0.44] |
| | ΔTI | -0.057 (0.123) | [-0.303, 0.179] | -0.011 (0.123) | [-0.252, 0.236] |
| **Theta** | TI | -0.156 (0.13) | [-0.413, 0.094] | 0.076 (0.113) | [-0.14, 0.3] |
| | ΔTI | -0.113 (0.13) | [-0.38, 0.132] | -0.013 (0.113) | [-0.234, 0.203] |
| **Alpha** | TI | 0.186 (0.116) | [-0.046, 0.414] | 0.07 (0.108) | [-0.137, 0.28] |
| | ΔTI | 0.071 (0.115) | [-0.163, 0.291] | -0.041 (0.107) | [-0.252, 0.169] |
| **Low-beta** | TI | -0.087 (0.141) | [-0.357, 0.198] | -0.085 (0.119) | [-0.317, 0.147] |
| | ΔTI | 0.117 (0.14) | [-0.169, 0.374] | 0.064 (0.12) | [-0.177, 0.294] |
| **High-beta** | TI | 0.202 (0.133) | [-0.057, 0.461] | 0.093 (0.114) | [-0.127, 0.317] |
| | ΔTI | **0.256 (0.131)** | **[0.003, 0.516]** | **0.252 (0.113)** | **[0.036, 0.48]** |
| ***Junior student sample*** | | | | | |
| **Broadband** | TI | **0.628 (0.279)** | **[0.047, 1.142]** | **0.369 (0.151)** | **[0.077, 0.666]** |
| | ΔTI | **0.52 (0.274)** | **[0.004, 1.063]** | 0.236 (0.153) | [-0.052, 0.545] |
| **Delta** | TI | 0.4 (0.259) | [-0.12, 0.885] | **0.389 (0.173)** | **[0.056, 0.729]** |
| | ΔTI | 0.279 (0.255) | [-0.224, 0.786] | 0.016 (0.172) | [-0.315, 0.354] |
| **Theta** | TI | -0.018 (0.265) | [-0.549, 0.492] | 0.147 (0.162) | [-0.19, 0.45] |
| | ΔTI | 0.157 (0.255) | [-0.346, 0.653] | 0.027 (0.161) | [-0.285, 0.344] |

| | | | | | |
|---|---|---|---|---|---|
| **Alpha** | TI | 0.319 (0.199) | [-0.098, 0.694] | 0.184 (0.173) | [-0.153, 0.524] |
| | ΔTI | -0.173 (0.188) | [-0.539, 0.206] | 0.093 (0.177) | [-0.247, 0.449] |
| **Low-beta** | TI | 0.067 (0.253) | [-0.423, 0.56] | 0.199 (0.204) | [-0.197, 0.594] |
| | ΔTI | 0.211 (0.243) | [-0.269, 0.692] | 0.271 (0.2) | [-0.112, 0.666] |
| **High-beta** | TI | **0.797 (0.246)** | **[0.297, 1.276]** | 0.203 (0.145) | [-0.091, 0.47] |
| | ΔTI | **0.643 (0.242)** | **[0.163, 1.114]** | **0.296 (0.148)** | **[0.006, 0.584]** |
| ***Senior student sample*** | | | | | |
| **Broadband** | TI | -0.128 (0.161) | [-0.452, 0.172] | -0.025 (0.152) | [-0.31, 0.281] |
| | ΔTI | -0.201 (0.161) | [-0.518, 0.115] | -0.059 (0.153) | [-0.353, 0.244] |
| **Delta** | TI | 0.12 (0.143) | [-0.148, 0.41] | 0.091 (0.165) | [-0.225, 0.412] |
| | ΔTI | -0.143 (0.141) | [-0.42, 0.134] | -0.027 (0.163) | [-0.368, 0.277] |
| **Theta** | TI | -0.211 (0.163) | [-0.535, 0.108] | 0.004 (0.151) | [-0.289, 0.293] |
| | ΔTI | -0.218 (0.163) | [-0.531, 0.103] | -0.039 (0.15) | [-0.327, 0.249] |
| **Alpha** | TI | 0.165 (0.156) | [-0.137, 0.476] | 0.051 (0.134) | [-0.211, 0.319] |
| | ΔTI | 0.191 (0.154) | [-0.116, 0.495] | -0.075 (0.135) | [-0.328, 0.201] |
| **Low-beta** | TI | -0.151 (0.179) | [-0.502, 0.2] | -0.174 (0.145) | [-0.454, 0.11] |
| | ΔTI | 0.104 (0.178) | [-0.248, 0.442] | -0.006 (0.146) | [-0.293, 0.279] |
| **High-beta** | TI | 0.057 (0.16) | [-0.272, 0.361] | 0.0 (0.168) | [-0.329, 0.335] |
| | ΔTI | 0.215 (0.16) | [-0.096, 0.537] | 0.213 (0.166) | [-0.113, 0.536] |

**Table S3.** Parametric estimation of static model results for score based on student-teacher coupling. For the full sample, N = 108 for Chinese and N = 129 for math. For the junior high school student sample, N = 37 for Chinese and N = 54 for math. For the senior high school student sample, N = 71 for Chinese and N = 75 for math.

| | | **Chinese Score** | **Wald chi2 (1)** | **Math Score** | **Wald chi2 (1)** |
|---|---|---|---|---|---|
| ***All student sample*** | | | | | |
| **Broadband** | TI | -0.119<br>(0.125) | 0.99 | -0.013<br>(0.134) | 0.01 |
| **Delta** | TI | 0.015<br>(0.125) | 0.02 | -0.220<br>(0.133) | 2.77 |
| **Theta** | TI | -0.046<br>(0.135) | 0.12 | **-0.237***<br>**(0.112)** | **4.64** |
| **Alpha** | TI | -0.129<br>(0.143) | 0.84 | -0.029<br>(0.140) | 0.04 |
| **Low-beta** | TI | 0.188<br>(0.134) | 2.06 | -0.112<br>(0.157) | 0.54 |
| **High-beta** | TI | 0.116<br>(0.146) | 0.65 | 0.159<br>(0.096) | 2.77 |
| ***Junior high scholl student sample*** | | | | | |
| **Broadband** | TI | 0.078<br>(0.139) | 0.39 | 0.262<br>(0.169) | 2.56 |
| **Delta** | TI | 0.108<br>(0.165) | 0.51 | -0.077<br>(0.146) | 0.30 |
| **Theta** | TI | 0.149<br>(0.129) | 1.44 | -0.001<br>(0.142) | 0.00 |
| **Alpha** | TI | -0.122<br>(0.136) | 0.86 | 0.029<br>(0.133) | 0.05 |
| **Low-beta** | TI | 0.108<br>(0.140) | 0.62 | 0.203<br>(0.139) | 2.31 |
| **High-beta** | TI | 0.029<br>(0.136) | 0.05 | 0.168<br>(0.127) | 1.79 |
| ***Senior high scholl student sample*** | | | | | |
| **Broadband** | TI | -0.237<br>(0.171) | 2.19 | -0.290<br>(0.263) | 1.22 |
| **Delta** | TI | 0.009<br>(0.147) | 0.00 | **-0.327***<br>**(0.160)** | **4.15** |
| **Theta** | TI | -0.196<br>(0.198) | 1.02 | **-0.515****<br>**(0.168)** | **9.35** |
| **Alpha** | TI | -0.061<br>(0.175) | 0.13 | -0.146<br>(0.196) | 0.55 |
| **Low-beta** | TI | 0.212<br>(0.155) | 2.15 | **-0.472***<br>**(0.215)** | **4.80** |

| | | | | | |
|---|---|---|---|---|---|
| **High-beta** | TI | 0.123 (0.218) | 0.34 | 0.261 (0.206) | 1.60 |

**Table S4.** Bayesian estimation of static model results for score based on student-teacher coupling. For the full sample, N = 108 for Chinese and N = 129 for math. For the junior high school student sample, N = 37 for Chinese and N = 54 for math. For the senior high school student sample, N = 71 for Chinese and N = 75 for math.

| | | **Chinese Score** | | **Math Score** | |
|---|---|---|---|---|---|
| | | **Posterior mean** | **HPDI 95%** | **Posterior mean** | **HPDI 95%** |
| ***All student sample*** | | | | | |
| **Broadband** | TI | -0.07 (0.081) | [-0.225, 0.094] | -0.004 (0.066) | [-0.137, 0.116] |
| **Delta** | TI | 0.014 (0.088) | [-0.16, 0.184] | -0.113 (0.07) | [-0.249, 0.025] |
| **Theta** | TI | -0.028 (0.082) | [-0.192, 0.128] | **-0.134 (0.064)** | **[-0.26, -0.013]** |
| **Alpha** | TI | -0.078 (0.087) | [-0.253, 0.087] | -0.015 (0.073) | [-0.164, 0.12] |
| **Low-beta** | TI | 0.109 (0.082) | [-0.051, 0.266] | -0.05 (0.069) | [-0.186, 0.082] |
| **High-beta** | TI | 0.064 (0.084) | [-0.101, 0.232] | 0.111 (0.067) | [-0.019, 0.245] |
| ***Junior student sample*** | | | | | |
| **Broadband** | TI | 0.075 (0.12) | [-0.157, 0.312] | 0.166 (0.111) | [-0.052, 0.379] |
| **Delta** | TI | 0.09 (0.126) | [-0.15, 0.341] | -0.062 (0.12) | [-0.291, 0.176] |
| **Theta** | TI | 0.11 (0.102) | [-0.087, 0.313] | -0.002 (0.105) | [-0.212, 0.206] |
| **Alpha** | TI | -0.091 (0.113) | [-0.3, 0.143] | 0.025 (0.118) | [-0.217, 0.248] |
| **Low-beta** | TI | 0.079 (0.115) | [-0.146, 0.311] | 0.149 (0.107) | [-0.06, 0.361] |
| **High-beta** | TI | 0.033 (0.114) | [-0.187, 0.264] | 0.141 (0.113) | [-0.071, 0.372] |
| ***Senior student sample*** | | | | | |
| **Broadband** | TI | -0.148 (0.115) | [-0.367, 0.082] | -0.105 (0.107) | [-0.317, 0.101] |
| **Delta** | TI | 0.005 (0.119) | [-0.23, 0.231] | **-0.194 (0.096)** | **[-0.382, -0.007]** |
| **Theta** | TI | -0.113 (0.118) | [-0.342, 0.121] | **-0.283 (0.097)** | **[-0.473, -0.088]** |

| | | | | | |
|---|---|---|---|---|---|
| **Alpha** | TI | -0.044 (0.119) | [-0.284, 0.185] | -0.075 (0.107) | [-0.284, 0.136] |
| **Low-beta** | TI | 0.152 (0.119) | [-0.078, 0.384] | **-0.217 (0.106)** | **[-0.421, -0.008]** |
| **High-beta** | TI | 0.068 (0.125) | [-0.182, 0.307] | 0.124 (0.102) | [-0.078, 0.323] |
